# Activity-driven synaptic translocation of LGI1 controls excitatory neurotransmission

**DOI:** 10.1101/2022.07.03.498586

**Authors:** Ulku Cuhadar, Lorenzo Calzado-Reyes, Carlos Pascual-Caro, Aman S. Aberra, Andreas Ritzau-Jost, Abhi Aggarwal, Keiji Ibata, Kaspar Podgorski, Michisuke Yuzaki, Christian Geis, Stefan Hallerman, Michael B. Hoppa, Jaime de Juan-Sanz

**Author notes:** These authors contributed equally to this work (Ulku Cuhadar, Lorenzo Calzado-Reyes).

## Abstract

The fine control of synaptic function requires robust trans-synaptic molecular interactions. However, it remains poorly understood how the landscape of trans-synaptic bridges dynamically remodels to reflect functional states of the synapse. Here we developed novel optical tools to visualize in firing synapses the molecular behavior of a particular secreted trans-synaptic protein, LGI1, and its presynaptic receptor, ADAM23, and discovered that neuronal activity acutely rearranges the abundance of these proteins at the synaptic cleft. Surprisingly, LGI1 in synapses was not secreted, as described elsewhere, but exo- and endocytosed through its interaction with ADAM23. Activity-driven translocation of LGI1 facilitated the formation of trans-synaptic connections proportionally to the history of activity of the synapse, adjusting excitatory transmission to synaptic firing rates. Thus, our findings reveal that LGI1 abundance at the synaptic cleft can be acutely remodeled and serves as a critical point of activity-dependent control of synaptic function.

**Highlights:**

– Neuronal activity translocates LGI1 and ADAM23 to the presynaptic surface
– LGI1 and ADAM23 are not located in synaptic vesicles
– Stable cleft localization of LGI1 depends on the history of synaptic activity
– LGI1 abundance at the synaptic cleft controls glutamate release

## Introduction

Physiological function of the mammalian brain relies on the orchestrated activity of excitatory and inhibitory synapses in neural circuits to encode the complexity of information driving cognition and behavior. Neurotransmission requires not only that individual pre- and post-synaptic sites work properly, but also that they are accurately connected in space through well-defined trans-synaptic interactions^1–8^. The fine control of the trans-synaptic molecular architecture is essential to maintain and dynamically modulate synaptic efficiency^9^, and it is now well established that distortions of trans-synaptic signaling cause synaptic dysfunction in several neuropsychiatric disorders, including epilepsy^5^. However, understanding how the complex molecular profile of a well-functioning synaptic connection is established, maintained and dynamically adjusted over time, and how altering these processes results in diseased brain states remains an unsolved research challenge.

The function of a particular trans-synaptic complex, formed between Leucin-rich, glioma inactivated 1 (LGI1) and its receptors ADAM22 and ADAM23, is essential to sustain circuit function *in vivo*, as lack of LGI1 function causes epilepsies of both genetic and autoimmune aetiology. Genetic mutations in LGI1 cause Autosomal Dominant Lateral Temporal Lobe Epilepsy (ADLTE)^10^ and autoantibodies against LGI1 cause a form of limbic encephalitis (LE) associated with cognitive decline and seizures^11–14^. Interestingly, ablating expression of LGI1 only in excitatory neurons causes epilepsy in mice, while selectively removing it from inhibitory neurons does not^15^, suggesting an important presynaptic role for LGI1 in the regulation of excitatory transmission in the brain. At the excitatory synaptic cleft, LGI1 acts as the molecular connector between presynaptic ADAM23 and postsynaptic ADAM22 receptors, linking structurally and functionally pre- and postsynaptic sites^11,16^. At the presynaptic level, LGI1-ADAM23 has been proposed to facilitate the function of the potassium channel Kv1.1, modulating the shape of the presynaptic action potential to curb activity-driven Ca^2+^ entry and reduce glutamate release^17–20^. Congruently, loss of LGI1 globally leads to increased glutamate release^15,21^, which provides a hypothesis on how LGI1 dysfunction could cause epilepsy. However, at the postsynaptic site LGI1-ADAM22 interacts with PSD-95 to enhance AMPA and NMDA receptor function^22–24^. In contrast to the increase in presynaptic function, loss of LGI1 decreases postsynaptic processing of neurotransmission, which has led us to propose that LGI1 controls postsynaptic glutamate receptor function exclusively in inhibitory neurons^22^. In this alternative model, LGI1 dysfunction decreases excitatory activation of inhibition to cause epilepsy.

While several studies using *in vivo* models have made clear that loss of LGI1 causes increased brain excitation and epilepsy^15,23,25–27^, controversy remains on how LGI1 localization and function can control the physiology of excitatory function, in part due to the lack of powerful tools to study the molecular behavior of LGI1 in single synapses. To tackle this, we developed a novel optical tool, LGI1-pHluorin (LGI1-pH), which allows monitoring for the first time LGI1 surface localization and trafficking in live firing synapses. Using this tool, we found that neuronal activity drives LGI1 translocation at presynaptic terminals, leading to an increased accumulation of LGI1 at the synaptic cleft of firing synapses. Increasing synaptic surface LGI1 levels narrows the presynaptic action potential waveform (AP), which in turn reduces AP-driven Ca^2+^ entry and glutamate release in a synapse-specific manner, suggesting that activity-driven molecular rearrangement of LGI1 trans-synaptic bridges modulates excitatory transmission correlatively to synaptic firing rates. Moreover, impairing LGI1 function by the presence of pathological autoantibodies against LGI1 led to a decrease in LGI1-pH at the surface and a corresponding increase in glutamate release. These experiments reveal a critical role for neuronal activity in shaping the molecular architecture of trans-synaptic connections, framing future investigations into the molecular control of neurotransmission and brain excitability by LGI1 and other trans-synaptic molecules.

## Results

### Neuronal activity drives LGI1 translocation to the synaptic surface

Early studies identified LGI1 as a secreted protein by expressing it in heterologous cell lines and measuring its constitutive secretion into the culture media, showing for the first time that several pathological mutations inhibited LGI1 secretion in this system^28,29^. However, while heterologous cell secretion assays can identify how mutations affect cellular trafficking^27,30–33^, they fail to mimic neuronal activity patterns and subcellular signaling pathways in neurons. Alternatively, expression of His-Flag tagged LGI1 has been used to visualize LGI1 at the surface of fixed neurons^23^, but this method lacks the spatiotemporal resolution to visualize the molecular behavior of LGI1 at the synaptic cleft in living neurons. To circumvent these limitations and explore with higher precision how the presence of LGI1 at the synaptic cleft is regulated, we developed a novel optical tool that allows selective visualization of surface-localized LGI1 molecules in live synapses. We fused pHluorin, a pH-sensitive variant of GFP, to the c-terminal end of LGI1 (see STAR methods). The fluorescence of pHluorin is quenched by acidic pH, which is typically found in the lumen of intracellular compartments such as synaptic vesicles, dense-core vesicles, secretory granules or presynaptic endosomes. However, when pHluorin is exposed to the extracellular neutral pH, it becomes ∼100 times more fluorescent^34^ (Figure 1A). We expressed LGI1-pH in primary excitatory hippocampal neurons that were co-cultured with astrocytes, a system that optimizes the optical access to single synapses and allows robustly quantifying dynamic changes in fluorescence in individual nerve terminals^35–37^. Selective expression in excitatory neurons was achieved by using the CaMKII promoter, as described previously^38^. Co-expression of LGI1-pH together with the presynaptic marker synapsin-mRuby confirmed the expected preferential localization of LGI1-pH at the synaptic surface of live neurons, showing ∼40% more relative synapse fluorescence at rest than a construct that simply expresses pHluorin in the entire surface of the neuron (Supp. Figure S1A-B).

**Figure 1.**
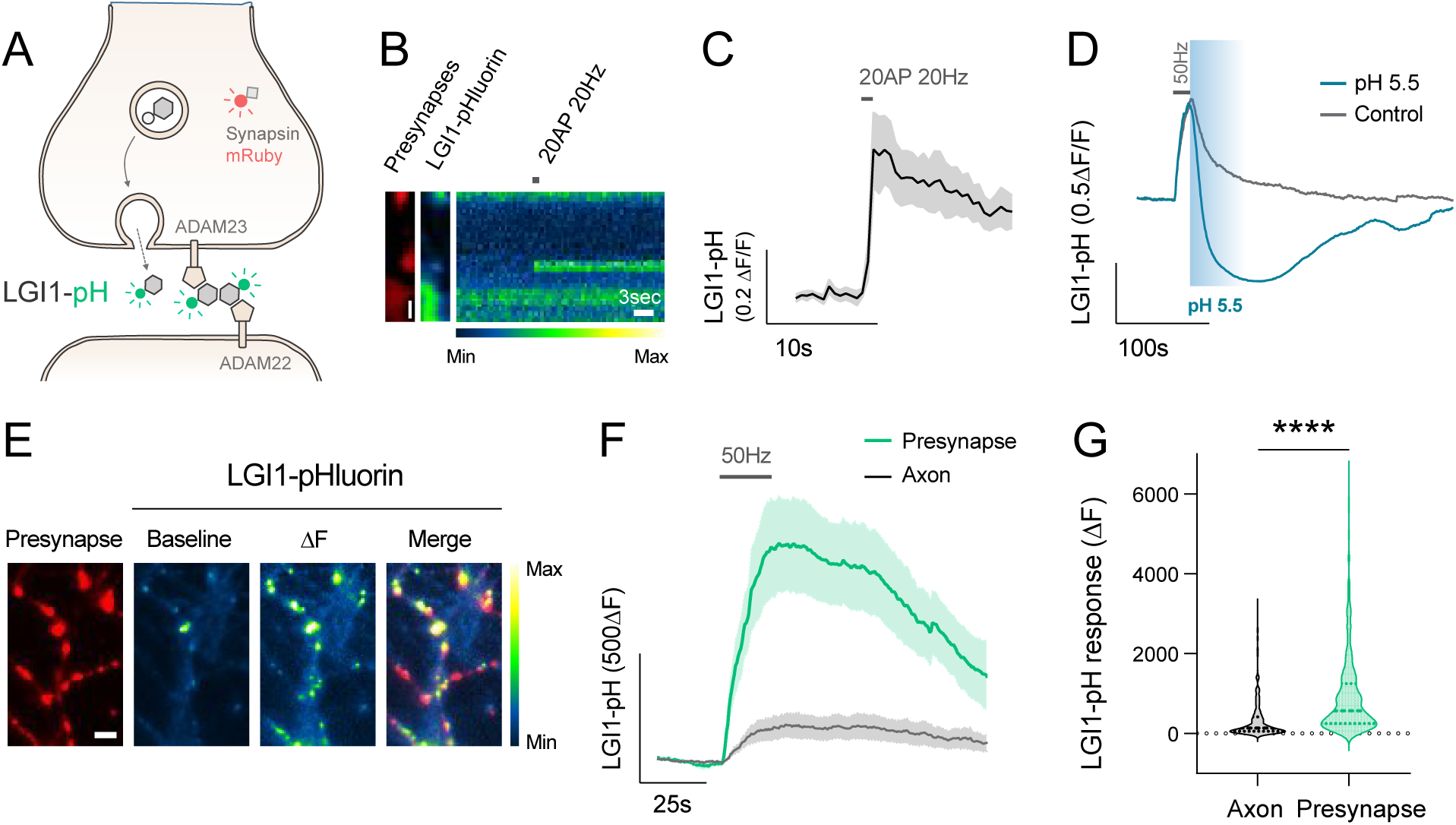
Neuronal activity drives LGI1 translocation to the synaptic surface. (**A**) Diagram showing LGI1-pHluorin (LGI1-pH), which is not fluorescent when located intracellularly (shown in grey) but becomes fluorescent when exposed at the surface (shown in green). Synapsin-mRuby is a presynaptic marker. **(B)** LGI1-pH signals in presynaptic boutons identified by synapsin-mRuby (left, colored in red) at rest (middle image) and during a 20AP 20Hz stimulus (displayed as a kymograph, right; pseudocolor scale shows low to high intensity). Time scale bar, 3s. Size bar, 2.4μm **(C)** LGI1-pHluorin response to 20AP 20Hz stimulus shown as average trace (N= 12 boutons, from 10 neurons, error bars are SEM). **(D)** Example traces of LGI1-pH response to 50Hz electrical stimulation during 20s in control or in the presence of pH 5.5 solution buffered with 2-(N-morpholino)ethanesulfonic acid (MES). **(E)** LGI1-pH response during 1000AP 50Hz electrical stimulation (ΔF) colocalizes with presynaptic bouton marker synapsin-mRuby; pseudocolor scale shows low to high intensity. Scale bar 4.8 μm. **(F)** Average traces of LGI1-pH responses to 1000AP 50Hz stimulation in synaptic boutons versus axonal regions of the same neurons (N= 5 neurons). Error bars are SEM. **(G)** Quantification of peak LGI1-pH response shown in F (n=5 neurons) from axonal (n=302) versus bouton regions (n=448). Mean ± SEM ΔF change is 346.2 ± 28.8 (a.u.) for axon and 937.8 ± 47.9 (a.u.) for synapses. Mann-Whitney test; ****p<0.0001).

We next quantified what fraction of LGI1 is present at the synaptic surface or inside presynaptic intracellular compartments by quenching the surface fraction using a non-permeable acidic solution and subsequently revealing the total amount of LGI1-pH using ammonium chloride (NH_4_Cl) (see STAR methods)^34^. Surprisingly, our estimates showed that ∼70% of synaptic LGI1 was located in presynaptic intracellular compartments (Supp. Figure S1C), an unexpectedly high fraction for a surface-localized neuronal protein^39^. We hypothesized that such a large internal pool ideally facilitates regulating the abundance of LGI1 at the synaptic surface on demand, permitting the translocation of a significant amount of LGI1 molecules during certain functional states, such as neuronal activity. To test this hypothesis, we electrically stimulated LGI1-pH-expressing neurons to mimic physiological firing paradigms of hippocampal place neurons in vivo, such as firing at 20Hz for 1 second^40^, and found translocation of LGI1 to the presynaptic neuronal surface in isolated boutons of the synaptic arborization (Figure 1B, C). We next verified that such presynaptic increase in LGI1-pH signal arose from surface accumulation. To facilitate LGI1 translocation in the majority of boutons, neurons were stimulated at 50Hz during 20 seconds and we subsequently confirmed that such increase was fully quenched by rapid perfusion of a low-pH solution (Figure 1D; see STAR methods). We noticed that LGI1-pH translocation to the surface occurred preferentially in synapses (Figure 1E). Quantifying changes in fluorescence in synapses versus inter-synapse axonal regions of the same axons revealed that virtually no change is found in the axon outside of presynaptic sites (Figure 1F, G). Conversely, no increase was observed in somas and dendrites as consequence of back-propagated action potentials, which contrarily generated a slight decrease in the signal (Supp. Figure S1D). Such decrease, however, is likely a reflection of activity-driven acidification of the endoplasmic reticulum, where LGI1-pH is transiently located while being synthetized, as an ER-localized pHluorin^36^ construct also reports a decrease in fluorescence during same stimulation paradigms (Supp. Figure S1E) and estimates of compartment pH obtained from neurons expressing ER-pHluorin or LGI1-pH in the dendrites appeared indistinguishable (Supp. Figure S1F). Taken together, these results show for the first time, to our knowledge, that neuronal activity controls LGI1 translocation to the synaptic surface at the presynapse.

### LGI1 is not located in synaptic vesicles

We next asked whether the intracellular pool of LGI1 is located in a structure different from synaptic vesicles (SVs) and examined this hypothesis with several complementary approaches. First, leveraging established methods to quantify pH inside organelles harboring pHluorin^34^, we found that the pH of the locale containing presynaptic LGI1 was around ∼6.1 in hippocampal neurons. This internal compartment, to which we will refer as LGI1 vesicles, presented a significantly more alkaline pH than our pH estimates from synaptic vesicles labeled with vGlut1-pHluorin (Figure 2A). Looking closely at the behavior of LGI1-pH in single boutons during electrical stimulation, we found that not all presynaptic sites responded simultaneously, but rather LGI1-pH exocytosis events appeared asynchronous and were delayed in many of the responding boutons (Figure 2B, lower panel). In contrast, SV exocytosis occurred in all boutons simultaneously (Figure 2B, top panel). We quantified the delay time between the stimulation start and the rise in fluorescence in individual boutons expressing either vGlut-pH or LGI1-pH, and found that LGI1-pH exocytosis events were delayed on average ∼4 seconds, being able to be delayed up to 20 seconds in some boutons (Figure 2C). Studying responses in individual boutons, we found that LGI1-pH exocytosis events presented dynamics compatible with univesicular exocytosis in at least one third of the recorded events, while the resting two thirds resembled dynamics of multivesicular exocytosis (Supp. Figure S2A, left panel; see STAR methods). A series of representative examples for each type of translocation, showing one, two or multiple exocytosis events are shown in Supp. Fig S2B-D, as well as the likelihood of each event type depending on the stimulation paradigm (Supp. Figure S2A). Stimulating for shorter periods of time mimicking physiological firing paradigms, such as firing at 20Hz for 1 second (Figure 1C), led uniquely to univesicular exocytosis (Supp. Figure S2A, right panel), suggesting that this may be the most likely mode of exocytosis occurring physiologically.

**Figure 2.**
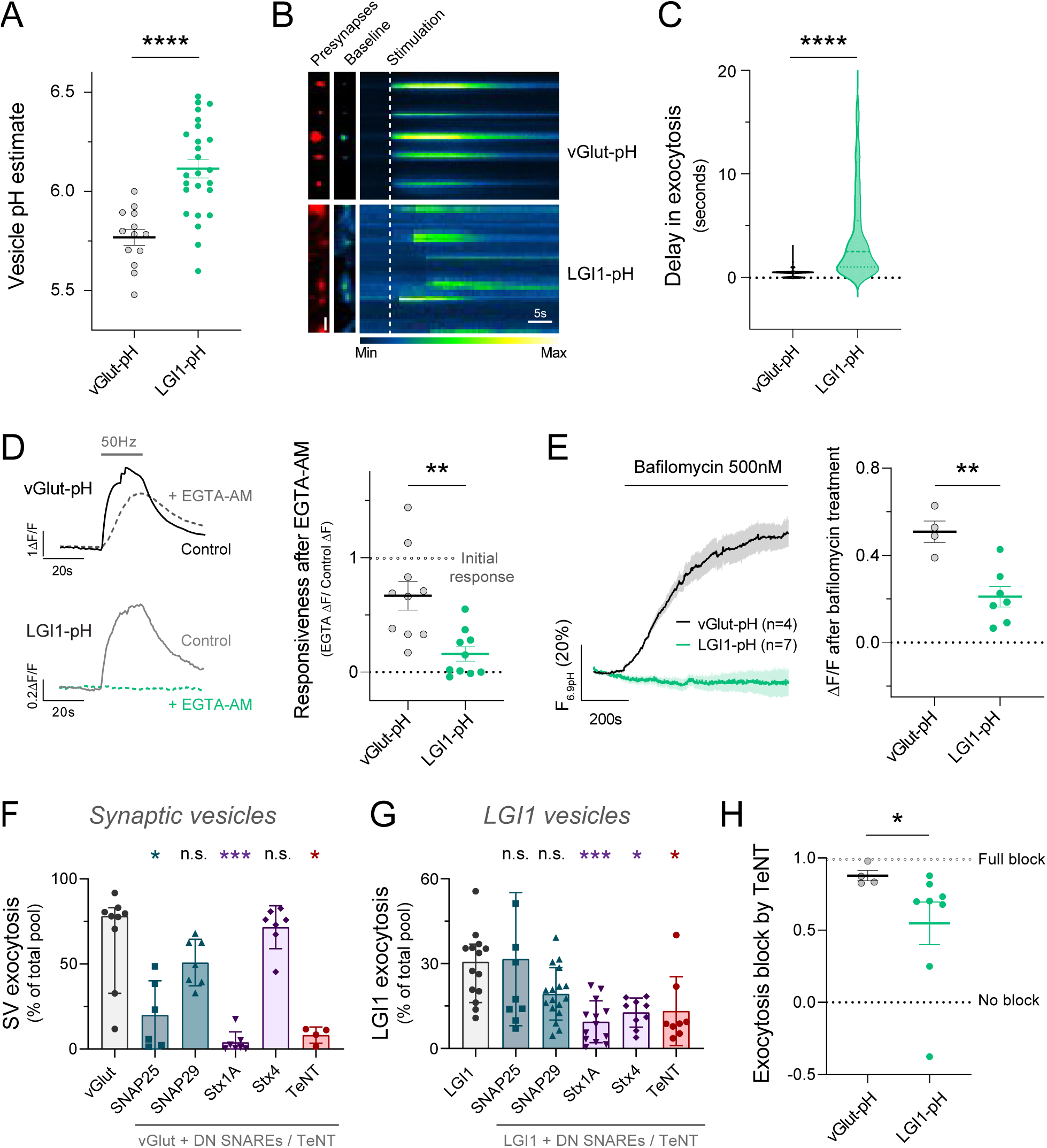
LGI1 is not located in synaptic vesicles. (**A**) Vesicle pH estimated using vGlut-pH (n=13 neurons) and LGI1-pH (n=25 neurons). Mean ± SEM vGlut-pH, 5.77 ± 0.04 pH units; LGI1-pH, 6.11 ± 0.05 pH units. t-test; ****p<0.0001. **(B)** vGlut-pH and LGI1-pH signals in boutons identified by synapsin-mRuby expression (left, colored in red) at rest (middle image) and during a 1000AP 50Hz stimulus (displayed as a kymograph, right; pseudocolor scale shows low to high intensity). Time scale bar, 5s. Size scale bar 2.4μm **(C)** Quantification of exocytosis delay in vGlut-pH (n=1214 boutons, n=11 neurons) versus LGI1-pH (n=390 boutons, n=7 neurons). Mann-Whitney test; ****p<0.0001. Mean ± SEM, 0.38 ± 0.01 seconds for vGlut-pH and 3.97 ± 0.22 seconds for LGI1-pH. **(D)** Left: Example traces of vGlut-pH (upper traces in black) and LGI1-pH (lower traces in blue) responses to 50Hz stimulation before (continuous line) and after (dotted line) 10 minutes of 2mM EGTA-AM treatment on the same neuron. Right: Quantification of the remaining response of vGlut-pH (n=10 neurons; 0.67 ± 0.12) and LGI1-pH (n=10 neurons; 0.16 ± 0.06) after EGTA-AM treatment. 1=no EGTA effect, 0=complete block by EGTA. t-test; **p<0.01. **(E)** Left: Responses to bafilomycin 500nM in vGlut-pH (n=4 neurons) and LGI1-pH (n=7 neurons). Changes are normalized to the maximal fluorescence change, obtained by applying NH_4_Cl at pH 6.9, the cytosolic pH. Right: quantification for vGlut-pH (mean ± SEM, 0.50 ± 0.05) and LGI1-pH (-0.04 ± 0.05) neurons. Mann-Whitney test; **p<0.01. **(F)** Synaptic vesicle exocytosis during 1000AP 50Hz in neurons expressing vGlut-pH with or without co-transfection of dominant-negative (DN) SNAREs or only vGlut-pH but treated for 24h with tetanus toxin. Data are normalized to maximum fluorescence change per bouton, obtained by applying NH_4_Cl at pH 7.4. (n=4-9 neurons). Dunn’s Multiple Comparison test; *p<0.05, ***p<0.001. **(G)** LGI1 exocytosis during 1000AP 50Hz in neurons expressing LGI1-pH with or without co-transfection of dominant-negative (DN) SNAREs or only LGI1-pH but treated for 24h with tetanus toxin. Data are normalized to maximum fluorescence change per bouton, obtained by applying NH_4_Cl at pH 7.4. (n=8-17 neurons). Dunn’s Multiple Comparison test; *p<0.05, ***p<0.001. **(H)** Exocytosis block by tetanus toxin (TeNT), calculated in neurons expressing vGlut-pH or LGI1-pH as 1-(TeNT response/average untreated response). Mann-Whitney test; *p<0.05.

Next, we reasoned that LGI1 exocytosis delays in individual boutons could be the consequence of a looser coupling between activity-driven Ca^2+^ entry and binding to the Ca^2+^ sensor present in LGI1 vesicles, making the process of exocytosis less efficient than in the case of synaptic vesicles, and thus, more prone to failure. To test this hypothesis, we incubated neurons expressing either vGlut-pH or LGI1-pH in the presence of EGTA-AM, a calcium chelator that impairs channel-vesicle coupling to greater extents in loosely coupled exocytosis mechanisms^41^. EGTA-AM exposure and concentration used had a relatively small impact on synaptic vesicle exocytosis during prolonged stimulation at 50Hz, allowing us to test whether in such conditions LGI1-pH exocytosis was significantly affected. Our results revealed that, contrarily to synaptic vesicles, LGI1-pH vesicle exocytosis was greatly blocked by the presence of EGTA-AM, thus revealing a much looser coupling to Ca^2+^ for exocytosis (Figure 2D).

We reasoned that if LGI1 vesicles are not synaptic vesicles, they should not contain neurotransmitter transporters. Recent data demonstrated that presence of neurotransmitter transporters in synaptic vesicles generates a steady-state H^+^ leak that is counteracted by the constitutive function of the vesicular H^+^-ATPase (vATPase), which continuously pumps H^+^ back to the synaptic vesicle lumen to maintain its H^+^ gradient^42^. Acutely blocking vATPase using bafilomycin reveals the H^+^ leak, which can be quantified using pHluorin (Figure 2E, black trace). As in SVs of excitatory neurons the H^+^ leak is driven by the presence of vGlut transporters^42^, we reasoned that if LGI1 vesicles are not synaptic vesicles, they should not present a H^+^ leak. We applied bafilomycin in neurons expressing LGI1-pH and confirmed that LGI1 vesicles, contrarily to synaptic vesicles, do not constitutively leak protons, indicating that they do not contain neurotransmitter transporters (Figure 2E).

Lastly, we reasoned that if LGI1 vesicles are distinct from synaptic vesicles, different SNARE proteins may control their exocytosis. We co-transfected different dominant negative SNAREs^43^ with either vGlut-pH or LGI1-pH and quantified cumulative exocytosis changes of single neurons during trains of action potentials. First, we confirmed that SNAP25 and Syntaxin-1A were required for synaptic vesicle exocytosis, as described elsewhere^44^, while SNAP29 and Syntaxin-4 did not impact this process, as expected^43^ (Figure 2F). Comparatively, lack of Syntaxin-4 function resulted in a significant reduction in LGI1-pH exocytosis during activity, which was blocked also by DN-Syntaxin-1A but not by DN-SNAP25 (Figure 2G). These results uncover a different mechanistic regulation between SV and LGI1 exocytosis. Previous work showed that presynaptic lysosomal exocytosis of Cerebellin-1, a secreted trans-synaptic protein, requires the function of Syntaxin-4 and SNAP29, and it is insensitive to tetanus toxin (TeNT), which cleaves VAMP1-3 proteins^43^. In contrast, we found that LGI1 exocytosis was not significantly impaired by the expression of DN-SNAP29 and, contrarily to Cerebellin-1, applying TeNT partially blocked LGI1 translocation (Figure 2G), suggesting that Cerebellin-1 and LGI1 do not rely on the same mechanisms for their presynaptic exocytosis. Interestingly, quantitative comparisons of the effect of TeNT in synaptic vesicles or LGI1 vesicles revealed that LGI1 exocytosis was not fully blocked (Figure 2H), suggesting that it may partially rely on TeNT-insensitive VAMP proteins for exocytosis. Taken together, these series of experiments demonstrate that LGI1 vesicles are not synaptic vesicles, despite being present at presynaptic sites and being able to undergo activity-driven exocytosis.

### LGI1 in neurons is primarily not secreted but trafficked bound to ADAM23

LGI1 is considered to be a secreted protein, and indeed, it can be found in the conditioned media of primary neurons^25^ and organotypic slices^45^. We leveraged the use of LGI1-pH to quantify to what extent the dynamics of activity-driven LGI1 exocytosis in single synapses resemble those of a canonical secreted protein, such as Neuropeptide Y (NPY)^46^. Field stimulation in neurons expressing NPY-pHluorin elicited a series of asynchronous exocytosis events whose dynamics showed a rapid increase and decrease in fluorescence (Supp. Figure S3A, B). Such fast decay in the signal after each exocytosis event is expected for a protein secreted into the medium, as it rapidly diffuses away from the secretion location, causing a rapid loss on fluorescence^46^. In contrast, we noted that individual LGI1-pH exocytosis events presented a much more sustained fluorescence over time after the initial increase (see examples at Supp. Figure S2B-D), indicating that molecules exocytosed stay at the secretion location for much longer times. To robustly quantify dynamics of LGI1-pH and NPY-pH univesicular exocytosis events, we aligned the temporal occurrence of asynchronous univesicular exocytosis responses from each protein to obtain a representative average of the dynamics of single univesicular exocytosis events (Figure 3A, B; see STAR methods). This analysis showed that LGI1-pH fluorescence remained on average mostly unchanged during at least 20 seconds after exocytosis in sharp contrast to NPY-pH, which by that time had already returned to baseline (Figure 3C). This contrasting result indicates that LGI1, on average, does not behave as a canonical secreted protein. Careful examination of 377 separate LGI1-pH uni- or multi-vesicular exocytosis events showed that, while none of the events presented dynamics resembling canonical secretion (as exemplified by NPY-pH), ∼8% events presented mixed kinetics with partial secretion-like decreases in fluorescence (Supp. Figure 3C, D; see STAR methods). The fast dynamics of these sharp decreases in LGI1-pH fluorescence at some point in the recording of single synapses is a readout that could be compatible with dissociation of LGI1-pH from the synaptic surface and its diffusion into the media. While the occurrence of these events was uncommon (29/377 boutons in 7 neurons), these data suggests that LGI1 can be secreted, or at least dissociated from the surface, and thus be present to some extent in the extracellular media. This is in agreement with previous work detecting endogenous LGI1 in the media of primary neuronal cultures^25^ and organotypic slices^45^. These results, however, unexpectedly suggest that when LGI1 is translocated to the neuronal surface it does not behave as a canonical secreted protein, as hypothesized elsewhere^11,47,48^.

**Figure 3.**
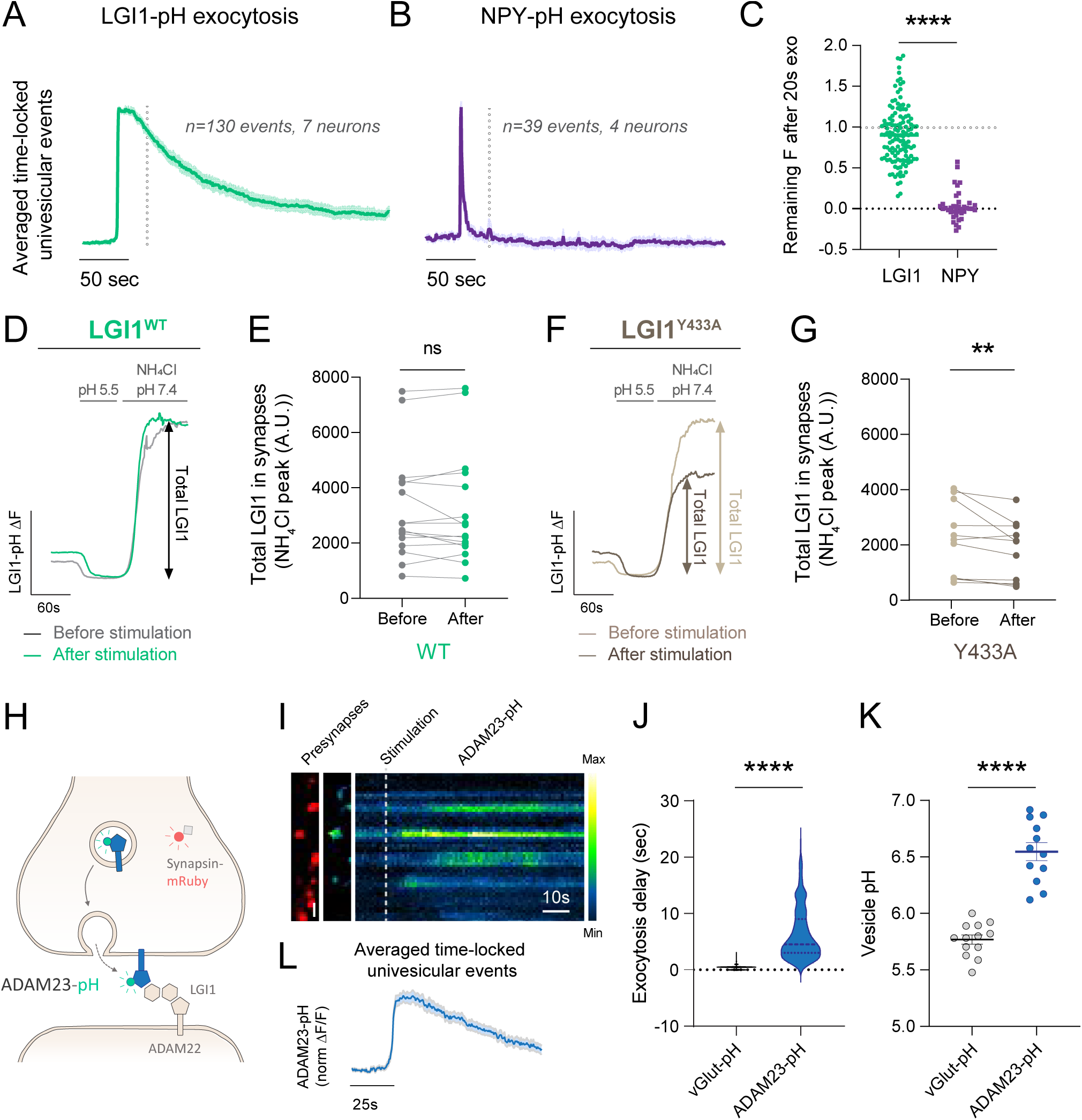
LGI1 is not secreted but exo and endo-cytosed bound to ADAM23. (**A**) Average of normalized univesicular exocytosis events from LGI1-pH, obtained by aligning temporally what otherwise would be asynchronous responses. Each single-bouton response was normalized to maximum fluorescence of the peak per event, averaging all the normalized events (n=130 events, 7 neurons). **(B)** Average of normalized univesicular exocytosis events from NPY-pH, obtained with the same analysis pipeline as in (A). (n=39 events, 4 neurons). **(C)** Quantification of the relative fluorescence in boutons expressing either LGI1-pH or NPY-pH after exocytosis (peak=1, baseline=0). Mann-Whitney test; ****p<0.0001. Mean ± SEM, 0.91 ± 0.03 for LGI1-pH and 0.03 ± 0.03 for NPY-pH. **(D)** Example traces of LGI1-pH responses to MES (pH 5.5) and NH_4_Cl (pH 7.4) before (gray) and after (green) electrical stimulation. **(E)** Quantification of total LGI1-pH levels before and after 1000AP 50Hz stimulation (n=16 neurons). Wilcoxon matched-pairs signed rank tests, n.s. **(F)** Example traces of LGI1^Y433A^-pH responses to Tyrode’s at pH 5.5 buffered with MES and NH_4_Cl (pH 7.4) before (light brown) and after (dark brown) electrical stimulation. **(G)** Quantification of total LGI1^Y433A^-pH levels before and after 1000AP 50Hz stimulation (n=11 neurons). Wilcoxon matched-pairs signed rank tests, **p<0.01. **(H)** Diagram showing ADAM23-pHluorin (ADAM23-pH), which is not fluorescent when located intracellularly (shown in grey) but becomes fluorescent when exposed at the surface (shown in green). **(I)** ADAM23-pHluorin signals in presynaptic boutons identified by synapsin-mRuby expression (left, colored in red) at rest (middle image) and during a 1000AP 50Hz stimulus (displayed as a kymograph, right; pseudocolor scale shows low to high intensity). Time scale bar 10s. Size scale bar 2.4μm. **(J)** Quantification of exocytosis delay in neurons expressing vGlut-pH (n=1214 boutons, n=11 neurons) or ADAM23-pH (n=257 boutons, n=6 neurons). Mean ± SEM, 0.38 ± 0.01 seconds for vGlut-pH and 6,34 ± 0,28 seconds for ADAM23-pH. Mann-Whitney test; ****p<0.0001. Note that the data shown for vGlut-pH is already shown in figure 2C but displayed here again for clarity when comparing it to ADAM23. **(K)** Vesicle pH in neurons expressing vGlut-pH (n=13 neurons) or ADAM23-pH (n=12 neurons). Mean ± SEM vGlut-pH; 5.77 ± 0.04; ADAM23-pH; 6.55 ± 0.08. t-test; ****p<0.0001. **(L)** Average dynamics of normalized univesicular exocytosis events from ADAM23-pH, obtained by aligning temporally what otherwise would be asynchronous responses. Each single-bouton response was normalized to maximum fluorescence of the peak per event, averaging all the normalized events (n=47 events, 6 neurons). Bars are SEM.

To quantitatively explore to what extent action potential firing may result in LGI1 secretion at the presynapse, we measured the total content of presynaptic LGI1-pH before and after stimulation. For these experiments, 1) we revealed the total pool of presynaptic LGI1 in the presence of NH_4_Cl in unstimulated neurons, 2) electrically stimulated them to expose ∼30% of the internal pool of LGI1-pH (as in Figure 2B) and 3) measured again the total pool of LGI1-pH at the same presynaptic sites. This paradigm should reveal the extent to which the internal pool was depleted by activity-driven secretion. These experiments did not reveal any detectable change in the total amount of LGI1-pH after stimulation (Figure 3D, E), suggesting that activity-driven surface translocation of LGI1-pH does not result in significant LGI1-pH secretion. These results also show that the total amount of LGI1-pH in a bouton remains constant despite translocation, thus making it unlikely that the gradual decay in LGI1-pH fluorescence observed after exocytosis is caused by LGI1-pH dissociation, which would result in detectable LGI1-pH loss. Contrarily, these results indicate that the reduction in fluorescence observed after exocytosis is likely caused by endocytosis and reacidification of recovered LGI1-pH molecules. These data further support the unexpected result that during activity LGI1 is generally not secreted but exo- and endo-cytosed.

Given that LGI1 does not contain a transmembrane domain and it can behave as a secreted protein in heterologous cells and neurons, we reasoned that its ability to undergo exo- and endo-cytosis has to be conferred through an interaction with a transmembrane receptor. At the presynaptic site, LGI1 interacts with ADAM23, a single-transmembrane receptor of the disintegrin and metalloproteinase domain-containing protein family^49^. Incorporating the Y433A mutation in LGI1 disrupts such interaction^16^, which would predict that LGI1^Y433A^-pH may behave as a secreted protein, as ADAM23 would not be able to retain it for subsequent endocytosis after translocation. To evaluate whether LGI1^Y433A^-pH is secreted during activity, we quantified the total pool of LGI1^Y433A^-pH in synapses before and after stimulation and found the total amount of LGI1^Y433A^-pH was significantly reduced after activity (Figure 3F, G). This suggests that if LGI1 cannot properly bind ADAM23, LGI1 molecules translocated during activity are more likely to be secreted and lost from the presynapse. Supporting this idea, careful examination of 477 events measured from 11 independent neurons expressing LGI1^Y433A^-pH identified that ∼15% events presented mixed kinetics with partial secretion-like decreases in fluorescence, roughly two-fold more than wild type LGI1-pH (Supp. Figure S3E).

We next hypothesized that if LGI1 requires ADAM23 for trafficking, ADAM23 should also undergo activity-driven translocation with similar dynamics. To test this, we designed a novel construct in which we cloned pHluorin after the signal peptide of the n-terminus of ADAM23, generating pHluorin-ADAM23 (ADAM23-pH, Figure 3H). Similarly to LGI1-pH, neuronal activity robustly drove asynchronous ADAM23-pH translocation on demand to the presynaptic surface (Figure 3I), presenting significant delays to undergo exocytosis, similarly to LGI1 (Figure 3J). Moreover, ADAM23-pH appeared to locate mostly in intracellular compartments at the presynapse that presented a pH more alkaline than SV pH (Figure 3K). Lastly, we aligned asynchronous univesicular ADAM23-pH exocytosis responses and obtained the representative average of the dynamics single univesicular exocytosis events, which presented similar kinetics to univesicular LGI1-pH exocytosis dynamics (Figure 3L). These results support the idea that during activity LGI1 is not secreted but undergoes exo- and endo-cytosis bound to ADAM23.

### Activity controls the stable localization of LGI1 at the synaptic surface

*In vivo* studies have shown that the extent to which LGI1 can be secreted, and thus presumably be present at the synapse, is a key controlling point for brain excitability^27^, yet the molecular control of LGI1 stabilization at the synaptic surface remains unknown. We noticed that in events undergoing univesicular exocytosis, fluorescence of LGI1-pH did on average not fully return to the baseline after endocytosis, suggesting that a fraction of new LGI1 molecules exposed were stabilized on the synaptic surface after activity (Figure 3A). This was also apparent when observing global LGI1-pH fluorescence changes in presynaptic arborizations of single neurons during stimulation (Figure 4A). This implies, however, that each exocytosis event should expose on average several LGI1-pH molecules, even in the cases of univesicular exocytosis. To confirm this, we used quantitative measurements of purified single EGFP molecules to estimate the number of LGI1-pH molecules exposed per univesicular exocytosis event. Leveraging the use of an EMCCD detector with single photon sensitivity and thermoelectric cooling to -95 °C to suppress Johnson–Nyquist noise (see STAR methods), we first imaged purified single EGFP molecules in a coverslip, which appeared as diffraction-limited spots (Supp. Figure S4A). Analysis of the intensity distribution of ∼1500 individual spots revealed a quantized distribution with a unitary size of ∼535 arbitrary units (Supp. Figure S4A), which we attributed to the fluorescence of a single EGFP molecule. Next, we imaged LGI1-pH using the same imaging conditions during action potential firing at 50Hz for 4 seconds and quantified changes in fluorescence in univesicular exocytosis events at single boutons (Supp. Figure S4B). Assuming that the brightness of EGFP and the bright state of pHluorin at pH 7.4 are equal^50,51^, exocytosis measurements can be calibrated in terms of the number of EGFP molecules. This estimate indicated that ∼6 LGI1-pH molecules are exposed to the synaptic surface when a single vesicle fuses (Supp. Figure S4C). While these shorter stimulations favored single univesicular exocytosis events (Supp. Figure S2A), we captured two subsequent exocytosis events in the same bouton in ∼12% of the cases (42 out of 331 events; Supp. Figure S2A; Supp. Figure S4D). We quantified the relative amplitude of the first and second exocytosis events and found that similar responses were obtained (median of population = 1.03; Supp. Figure S4E), indicating that the amount of LGI1-pH present in different LGI1 vesicles of the same bouton is on average constant. These results support the idea that each LGI1 vesicle may contain several LGI1 molecules, which in turn can enable synaptic stabilization of a fraction of the molecules translocated.

**Figure 4.**
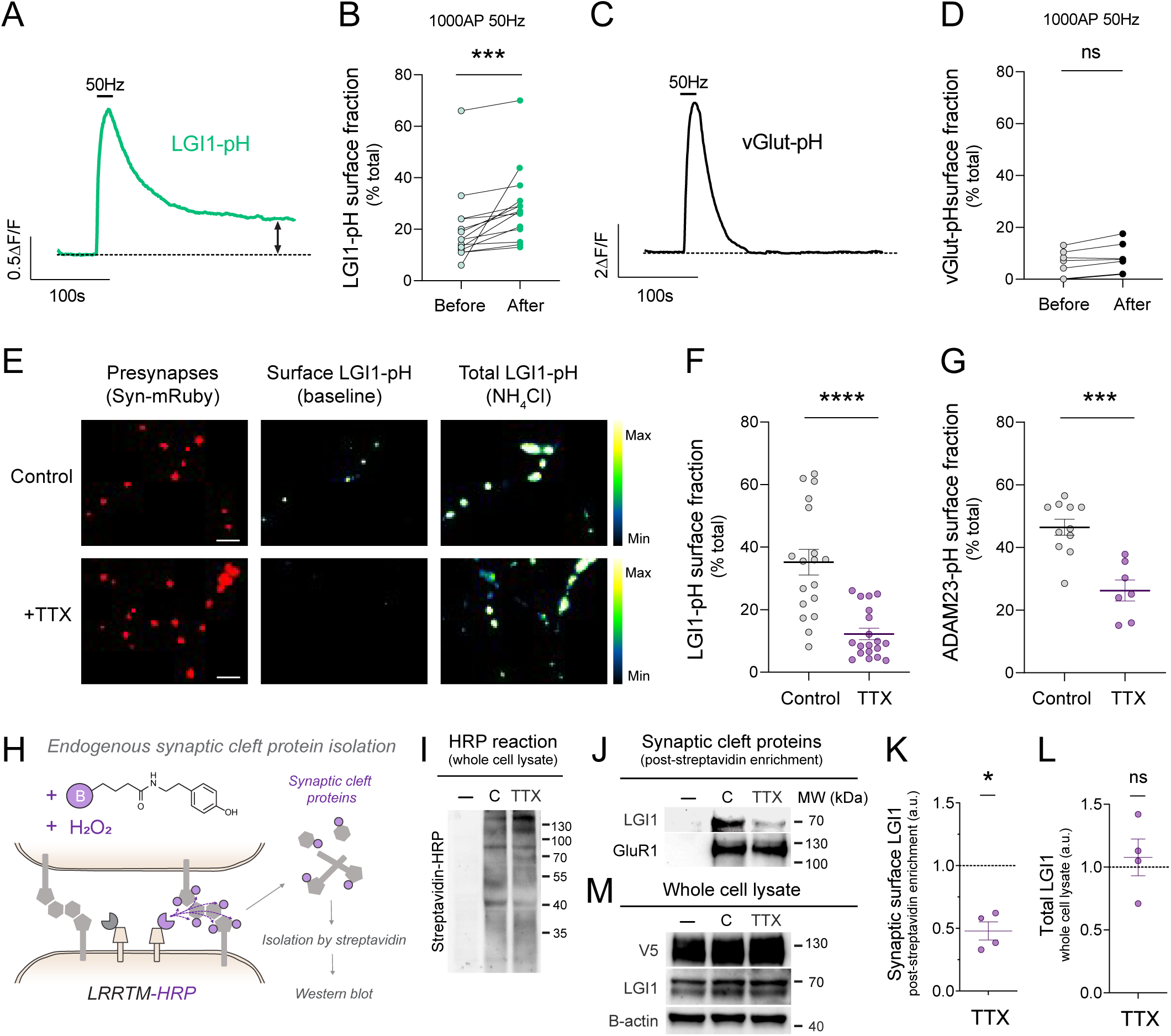
Stable localization of LGI1 at the cleft depends on the history of synaptic activity. (**A)** Example response of a single presynaptic arborization expressing LGI1-pH during 1000AP 50Hz electrical stimulation. Dotted line indicates the baseline before stimulation. **(B)** LGI1-pH fraction present at the synaptic surface before and after electrical stimulation (1000AP 50Hz) for individual neurons (n=14 neurons). Wilcoxon matched-pairs signed rank test; ***p<0.001. **(C)** Example response of a single presynaptic arborization expressing vGlut-pH during 1000AP 50Hz electrical stimulation. Dotted line indicates the baseline before stimulation. **(D)** vGlut-pH fraction present at the synaptic surface before and after electrical stimulation (1000AP 50Hz) for individual neurons (n=6 neurons). Wilcoxon matched-pairs signed rank test; n.s. **(E)** Example images of LGI1-pHluorin expression levels at the surface (middle panel) and total (right panel, revealed by NH4Cl pH 7.4) in control neurons versus neurons treated with TTX for 5 days. Presynaptic boutons are identified by synapsin-mRuby (left, in red). Pseudocolor scale shows low to high intensity. Scale bar 4.8μm **(F)** Fraction of LGI1-pH present at the synaptic surface in control (n=18 neurons) versus neurons treated with TTX for 5 days (n=19 neurons). Mean ± SEM, 35.2 ± 4.1 for control and 12.2 ± 1.8 for TTX. Mann-Whitney test, ****p<0.0001. **(G)** Fraction of ADAM23-pH present at the synaptic surface in control (n=11 neurons) versus TTX treated neurons (n=7 neurons). Data are mean ± SEM, 46.4 ± 2.5 for control and 26.3 ± 3.4 for TTX. Mann-Whitney test, ***p<0.001. **(H)** Diagram showing experimental flow to biochemically isolate endogenous synaptic cleft proteins (see STAR methods). **(I)** Example blot for total biotinylation rates in neurons expressing LRRTM-HRP treated with or without TTX for 5 days; Minus signal (-) indicates a negative control in which reaction was not run. **(J)** Example western blot experiment showing endogenous LGI1 levels isolated from the synaptic surface in control and neurons treated with TTX for 5 days (top panel), with the same experimental conditions shown in (H). As a control, endogenous levels of synaptically-localized GluR1 remained unchanged (lower panel). **(K)** Example western blot experiment showing total LGI1 levels in control and neurons treated with TTX for 5 days. V5 shows the expression of LRRTM-HRP-V5 and b-actin is a loading control. **(L)** Quantification of endogenous levels of LGI1 at the synaptic surface in TTX treated neurons, Mann-Whitney test, *p<0.05; **(M)** Quantification of endogenous total levels of LGI1 obtained from whole cell lysate of the experiments shown in (L). Mann-Whitney test, n.s.

To quantify precisely the extent to which LGI1-pH stabilizes at synapses after neuronal activity, we measured the surface fraction of LGI1-pH at the presynapse before and after stimulation (see STAR methods). While the total amount of LGI1 remains unchanged in the conditions tested (Figure 3D, E), the percentage of LGI1 molecules at the synaptic surface stably increased by ∼30% when measured 5 to 10 min after firing (Figure 4B). In a different set of experiments, we stimulated three times longer at the same frequency, which also increased LGI1-pH stabilization by ∼30% (Supp. Figure S5A), indicating that activity-driven increases in surface LGI1 may reach a saturation point, at least for a single stimulation train. As a control, the same stimulation paradigm did not induce surface stabilization of vGlut-pH, indicating that the activity-driven synaptic stabilization observed in LGI1 is not generalizable to other presynaptic proteins that traffic during firing (Figure 4C, D). These results indicate that the presence of LGI1 at the synaptic surface in a given time is controlled by the history of firing of such synapse.

To test this hypothesis, we reasoned that inhibiting spontaneous firing in culture for several days should decrease synaptic surface localization of LGI1. We transfected LGI1-pH and two days later neurons were treated for five days with tetrodotoxin (TTX), a selective inhibitor of neuronal Na^+^ channels that results in blockage of action potential propagation. We found that LGI1-pH was hardly detectable in the surface of TTX-treated neurons (Figure 4E), which presented on average a reduction in LGI1 surface localization of 70% compared to untreated neurons (Figure 4F). Similar experiments using ADAM23-pH showed the same phenotype, supporting the idea that ADAM23 and LGI1 pHluorin are trafficked together to the synaptic surface (Figure 4G). We confirmed that loss of these proteins at the surface was not a consequence of loss of expression induced by TTX, as both LGI1-pH and ADAM23-pH total pools are not reduced. In fact, we observed that LGI1-pH was even slightly increased (Supp. Figure S5B, C).

To confirm that history of activity controls surface localization of endogenously expressed LGI1, we isolated synaptic cleft proteins in control and TTX-treated neurons using recent technologies for synaptic cleft proximity biotinylation^52^ (Figure 4H). In this technique specific isolation of synaptic cleft proteins is achieved by expressing only in excitatory synapses a postsynaptic surface protein from the LRRTM family fused to a biotinylating enzyme (HRP), and running a local biotinylation reaction for 1 minute exclusively in the synaptic surface leveraging the use of a biotin-phenol variant that cannot cross the plasma membrane^52^ (Figure 4H). We first confirmed that TTX treatment did not impair synaptic cleft biotinylation rates (Figure 4I). In these conditions, TTX treated neurons presented a significant reduction in the abundance of endogenous LGI1 at the synaptic cleft (Figure 4J, K). As controls, the abundance in the synaptic surface of other synaptic cleft proteins, such as GluR1, remained unchanged, and the total amount of endogenous LGI1 was not affected by TTX treatment (Figure 4L, M). Taken together, these experiments show that neuronal activity is a major regulatory element in the control of LGI1 surface localization at excitatory synaptic clefts and show that the extent to which LGI1 is present at the synaptic surface reflects the history of activity of the synapse.

### Epilepsy-associated LGI1 mutants present translocation or stabilization defects at the synapse

Our novel tools have enabled us to discover previously unrecognized mechanisms controlling LGI1 abundance at the synaptic cleft. Loss of LGI1 function at the synaptic surface is thought to cause epilepsy because most pathogenic mutations inhibit LGI1 protein secretion in heterologous cells^10,25,27,30,31^. However, a few disease-causing LGI1 mutants can be easily found in the media of transfected heterologous cells^32^, and thus what is dysfunctional in these cases remains poorly understood. We reasoned that LGI1-pHluorin should enable straightforward quantitative analyses of the molecular dysfunction of LGI1 disease-causing mutations. We generated a series of LGI1-pHluorin mutants that in HEK cells are known to be secretion-defective (C200R, E383A) or secretion-positive (S473L, R474Q), together with a mutant (T380A) whose behavior remains controversial, as it has been claimed to be both secretion-defective^27^ and secretion-positive^32^. Mutations associated with epilepsy in humans are typically heterozygous and thus we studied LGI1-pH mutants in the presence of endogenous LGI1 to better understand how each mutant may misbehave at the synapse. We first saw that pHluorin variants of secretion-defective mutants C200R and E383A appeared mostly retained in the neuronal somatic ER, did not show any detectable surface LGI1 in synapses at rest (not shown) and did not undergo activity-driven translocation (Figure 5A, red-colored mutants). In contrast, mutant T380A showed a reduced, yet detectable presence at the synaptic surface (Supp. Figure S6A, B), and electrical stimulation drove its translocation to the synaptic cleft, although with much less efficiency than wild type LGI1 (Figure 5A, purple-colored). Mutants identified as secretion-positive in heterologous systems, such as S473L, Y433A and R474Q, were found at the synaptic surface in levels comparable to wild type LGI1 (Supp. Figure S6A) and their activity-driven translocation was indistinguishable from wild type LGI1 (Figure 5A).

**Figure 5.**
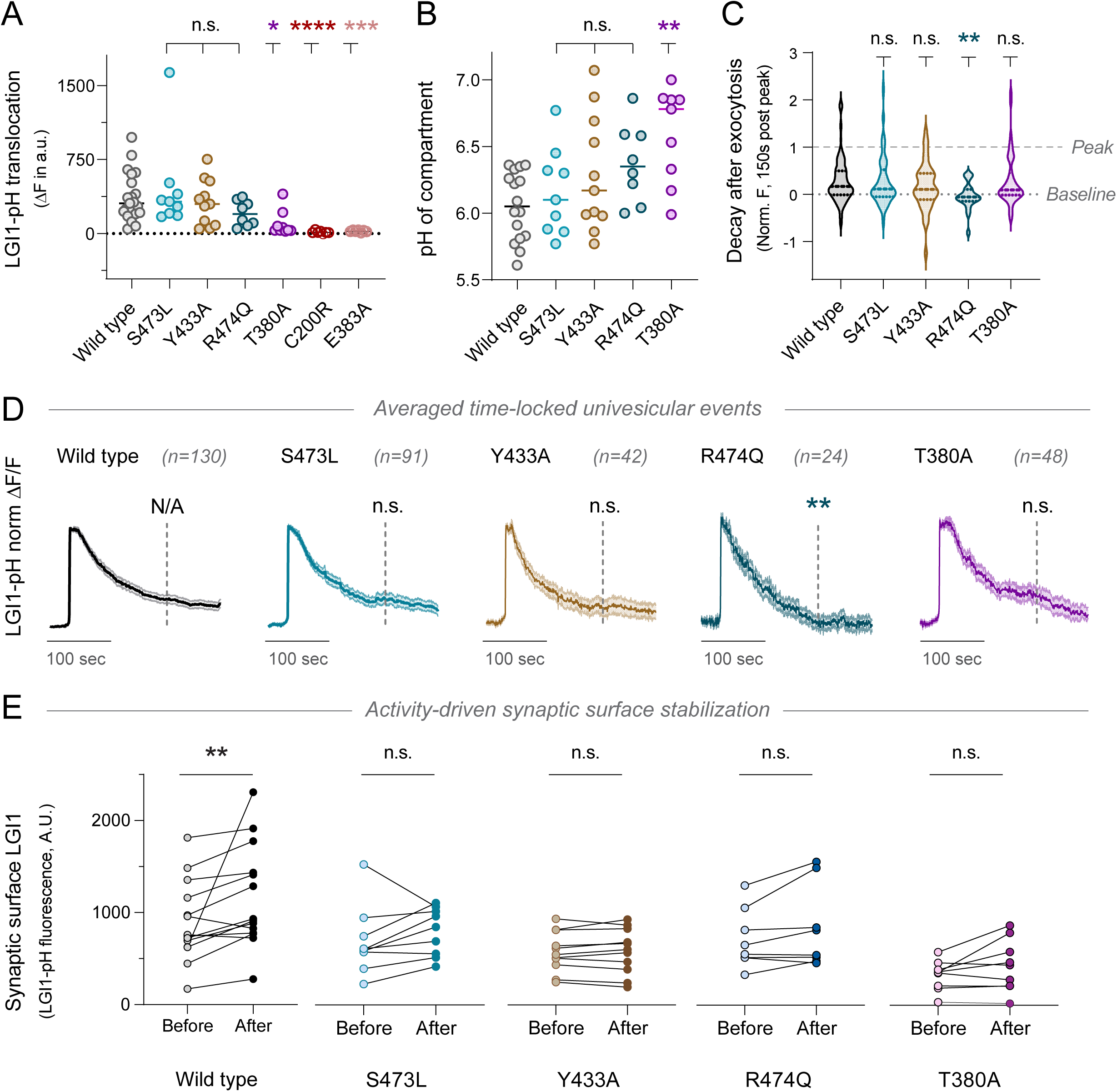
Dysfunctional translocation dynamics and stabilization of epilepsy-associated LGI1 mutants. (**A**) Average ΔF peak during 1000AP 50Hz stimulation of pHluorin-tagged LGI1 constructs WT, S473L, Y433A, R474Q, T380A, E383A and C200R. Data are mean ± SEM in a.u., WT, n=21, 366.5 ± 53.1; S473L, n=9, 448.6 ± 152.5; Y433A, n=11, 314.4 ± 69.8; R474Q, n=8, 205.7 ± 45.0; T380A, n=9, 99.9 ± 42.5; C200R, n=7, 13.5 ± 5.7; E383A, n=7, 23.4 ± 3.6. Dunn’s multiple comparisons test of the WT peak response vs. S473L,. Y433A or R474Q showed no statistical difference (n.s.); WT vs. T380A *p<0.05; WT vs. E383A ***p<0.001; WT vs. C200R ****p<0.0001. Note that data corresponding to WT LGI1 was already shown in Figure 3A. **(B)** pH of intracellular compartments containing pHluorin-tagged LGI1 constructs WT, S473L, Y433A, R474Q or T380A. Data are mean ± SEM in pH units; WT, n=16, 6.05 ± 0.06 pH units; S473L, n=9, 6.16 ± 0.11; Y433A, n=11, 6.11 ± 0.14; R474Q, n=8, 6.37 ± 0.10; T380A, n=9, 6.60 ± 0.12. Dunn’s multiple comparisons test of WT pH vs. S473L, Y433A or R474Q showed no statistical difference (n.s.); WT vs. T380A, ***p<0.001. **(C)** Quantification of fluorescence decay in individual boutons 150 seconds after exocytosis peak for LGI1-pH constructs WT, S473L, Y433A, R474Q or T380A. Data per bouton were normalized to maximum fluorescence of the peak response in that bouton. Data are mean ± SEM. WT, n= 130, 0.27 ± 0.04; S473L, n= 91, 0.27 ± 0.06; Y433A, n= 42, 0.17 ± 0.06; R474Q, n= 24, -0.03 ± 0.05; T380A, n= 48, 0.28 ± 0.08. Dunn’s multiple comparisons test of WT decay vs. S473L, Y433A or T380A showed no statistical difference (n.s.); WT vs. R474Q **p<0.01. **(D)** Average of normalized univesicular exocytosis events during electrical stimulation (1000AP 50Hz) of pHluorin-tagged LGI1 constructs WT, S473L, Y433A, R474Q and T380A, obtained by aligning temporally what otherwise would be asynchronous responses. Each single-bouton response per condition was normalized to maximum fluorescence of the peak per event, averaging all the normalized events for WT (n=130 events, 6 neurons), S473L (n=91 events, 9 neurons), Y433A (n=42 events, 9 neurons), R474Q (n=24 events, 8 neurons) and T380A (n=48 events, 9 neurons). Bars are SEM. Dashed lines in each trace indicate the time at which fluorescence was measured to estimate decay rates. Statistical tests on the dashed lines show the Dunn’s multiple comparisons test of the decay as shown in C. **(E)** Stable synaptic surface LGI1-pH change after 1000AP 50Hz electrical stimulation, measured 5-10 min after stimulation, for WT (n=14 neurons), S473L (n=9 neurons), Y433A (n=11 neurons) R474Q (n=8 neurons) and T380A (n=9 neurons). To evaluate whether each particular mutant can be stabilized or not at the synapse surface, each individual condition was tested separately through Wilcoxon matched-pairs signed rank test; **p<0.01.

We next measured the pH of the presynaptic intracellular compartment where different mutants were located at the presynapse. We found that LGI1^T380A^ was in a compartment significantly less acidic than wild type LGI1, with a pH ∼7 (Figure 5B). This is likely the consequence of this mutant being in LGI1 vesicles (pH ∼6.1) but also partially retained in axonal ER, which has a pH ∼7.2^36^, and thus the final measurement reflects a mix of both populations at the synapse. The partial localization of LGI1^T380A^ at the ER also matches with its detectable but reduced capacity to undergo activity-driven translocation, as ER-localized LGI1 molecules should in principle be unable to undergo exocytosis. Lastly, using NH_4_Cl at pH 7.4 to reveal all pHluorin molecules, we found a reduced number of LGI1^T380A^-pH molecules localized at the presynapse (Supp. Figure S6C). Taken together, these experiments suggest that LGI1^T380A^ suffers partial retention in the ER, which results in a reduced number of translocation-competent molecules at presynaptic sites. These data resolve the current controversy around the molecular behavior of this mutant^27,32^ by demonstrating that its translocation is severely impaired, yet detectable. We confirmed that secretion-competent S473L, Y433A and R474Q mutants appeared to be expressed in similar levels to wild type LGI1 at presynapses (Supp. Figure S6C), in agreement with their strong capacity to be translocated to the surface during activity (Figure 5A).

Exocytosis of S473L, Y433A, R474Q and T380A occurred asynchronously in different boutons of the same neuron as expected (not shown) and thus we again time-locked univesicular events to quantitatively compare translocation dynamics of the different mutants without the contribution of asynchronicity. No distinguishable differences were observed in the short-term decay of fluorescence between the wild type and the S473L, Y433A, or T380A variants, measured by quantifying the remaining signal 150 seconds (Figure 5C, D) or 210 seconds after peak responses (Supp. Figure S6D). However, we found that the translocation-competent R474Q mutant decayed faster after exocytosis (Figure 5C, D; Supp. Figure S6D). The R474Q mutation causes both a block of the LGI1-LGI1 interaction^16^ and a reduction in LGI1’s affinity for ADAM22^53^, disrupting the higher-order assembly of the LGI1–ADAM22/23 complex. It is thus possible that the different translocation dynamics of LGI1^R474Q^ reflect a much reduced probability of establishing new LGI1–ADAM22/23 complexes during synaptic exposure of LGI1^R474Q^, presenting less likelihood of synapse stabilization and thus faster short term retrieval. Interestingly, mutations that block the interaction with ADAM23 (Y433A)^16^ and ADAM22 (Y433A, S473L)^27^ without impairing LGI1-LGI1 interactions, presented translocation dynamics indistinguishable from wild type LGI1 (Figure 5C, D). It is possible that this is a consequence of the preserved interaction between LGI1-pH mutants and endogenously expressed wild type LGI1, which itself binds ADAM22/23 receptors.

As activity drives long term stabilization of wild type LGI1 at the presynaptic surface (Figure 4A, B), we analyzed whether LGI1 mutants presented a similar phenotype. Individual testing of activity-driven LGI1 stabilization per mutant revealed that none were significantly increased at the synaptic surface after 5-10 min of neuronal activity (Figure 5E), in contrast to wild type LGI1 (Figure 4A, B; Figure 5E). This inability to be stabilized at the cleft surface likely arised from a reduced capacity to bind ADAM22 at the postsynapse, which has been shown in S473L^27^, Y433A^16^, R474Q^53^ and T380A^32^ LGI1 mutants. On the other hand, as we found that losing ADAM23 binding resulted in activity-driven secretion of LGI1^Y433A^-pH (Figure 3F, G), we explored whether other mutants presented this phenotype. We found that neither S473L, R474Q or T380A presented detectable loss on the total synaptic protein levels after neuronal activity, which we measured by revealing the total pool of LGI1-pH mutants in the presence of NH_4_Cl at pH 7.4 before and after 1000AP 50Hz stimulation (Supp. Figure S6E). Binding to ADAM23 has been shown to be preserved for S473L^27,53^ and R474Q^53^, and thus this further supports the idea that losing interaction with ADAM23 may lead to LGI1 secretion (Figure 3F, G), but if that interaction is preserved, LGI1 is recovered back and no total protein is lost (Supp. Figure S6E, F). However, we did not detect secretion for LGI1^T380A^, even though it has been reported to be unable to interact with ADAM23^32^. This result, however, is complex to interpret because during stimulation significantly fewer LGI1-pH molecules are translocated (Figure 5A) and the theoretical capacity for secretion, and thus the ability of detecting it, is significantly reduced (Supp. Figure S6G). Additionally, LGI1^T380A^-pH likely presents a mixed intracellular localization in both axonal ER and LGI1 vesicles (Figure 5B), and applying NH_4_Cl at pH 7.4 reveals molecules at both compartments, resulting in a contaminated measurement that does not allow measuring LGI1 loss from translocating vesicles exclusively.

Given that LGI1 modulates the function of the potassium channel Kv1.1^18–20,54^, overexpressing different mutants in these experiments could impact action potential invasion of axonal boutons differently in each condition, potentially inducing failures of action potential propagation that could drive differences in exocytosis recruitment for each mutant. To control for this, we fused wild type LGI1 and mutants to pHmScarlet^55^, a recently developed red pH-sensitive fluorescent protein, and measured possible action potential failures in axonal boutons using jGCaMP8f, the last generation of fast and sensitive genetically encoded Ca^2+^ sensors^56^. Stimulating neurons to fire one action potential every second during 50 seconds, we found that action potentials propagated equally well in untransfected neurons or neurons expressing wild type LGI1, secretion-competent (R474Q, T380A) or secretion-defective (E383A) mutants (Supp. Figure S7A, B). This indicates that differences in exocytosis in each case are not a consequence of differential failures in action potential propagation. Taken together, these experiments demonstrate that our novel tools provide the field with a powerful approach to dissect the pathogenicity of newly-identified LGI1 mutations, which should help define whether novel variants identified by molecular genetic testing could indeed cause epilepsy.

### LGI1 surface abundance in individual boutons serves as a major modulatory mechanism of glutamate release

Loss of LGI1 increases glutamate release from excitatory hippocampal neurons^21,57^, as LGI1 constitutively constrains the function of the potassium channel Kv1.1, curbing activity-driven Ca^2+^ entry and reducing glutamate release^18–20,54^. Thus, we reasoned that the extent to which LGI1 is present at the surface of individual boutons should result in proportional synapse-specific readjustments of the magnitude of AP-driven Ca^2+^ entry and glutamate release. To test this hypothesis, we used LGI1-pHmScarlet to combine single bouton measurements of LGI1 surface localization with activity-driven Ca^2+^ entry or glutamate release (Supp. Figure S8A). Synaptic surface LGI1-pHmScarlet was reduced by TTX (Supp. Figure S8B), as observed for LGI1-pH (Figure 4E, F) and endogenous LGI1 (Figure 4J, K), indicating that LGI1-pHmScarlet fluorescence reflects LGI1 surface localization. We first co-expressed LGI1-pHmScarlet with either jGCaMP8f^56^ or iGluSnFR3^58^ to measure presynaptic Ca^2+^ or glutamate release in TTX-treated neurons and found that they presented a ∼30% increase in Ca^2+^ entry (Supp. Figure S8C-E) and a ∼50% increase in glutamate release (Supp. Figure S8F-H). These initial results provided phenotypes compatible with the idea that surface LGI1 abundance may control synaptic function. However, these results did not prove a causal relationship between surface LGI1 levels and synaptic function modulation, as chronic TTX treatment is known to modulate additional signaling pathways *a priori* unrelated to LGI1 that can also control presynaptic strength^59^.

We thus next leveraged the use of an optical voltage sensor, Archon1^60^, which robustly replicates voltage changes measured by electrophysiology in patched somas of neurons^61^, to quantify directly whether increasing the presence of LGI1 at synapses could modulate locally Kv function. We found that overexpressing LGI1-pHmScarlet resulted in a significantly narrowed presynaptic action potential waveform (Figure 6A), particularly in the end phase of repolarization (Supplementary Table ST1). In contrast, the amplitude of the AP remained unaltered (Figure 6B). Such modulation suggests differences in Kv1 function, as blockade of Kv1.1/1.2 channels broadens the AP waveform at the presynapse^62,63^. Given that the most significant modulator of calcium entry during an action potential is the end phase of repolarization^64^, we reasoned that this specific narrowing of the AP waveform would curb both AP-driven Ca^2+^ entry and glutamate release. We overexpressed LGI1-pHmScarlet and found that increasing LGI1 levels decreased both presynaptic Ca^2+^ entry (Figure 6D, E) and glutamate release (Figure 6G, H) by 25% and 40%, respectively.

**Figure 6.**
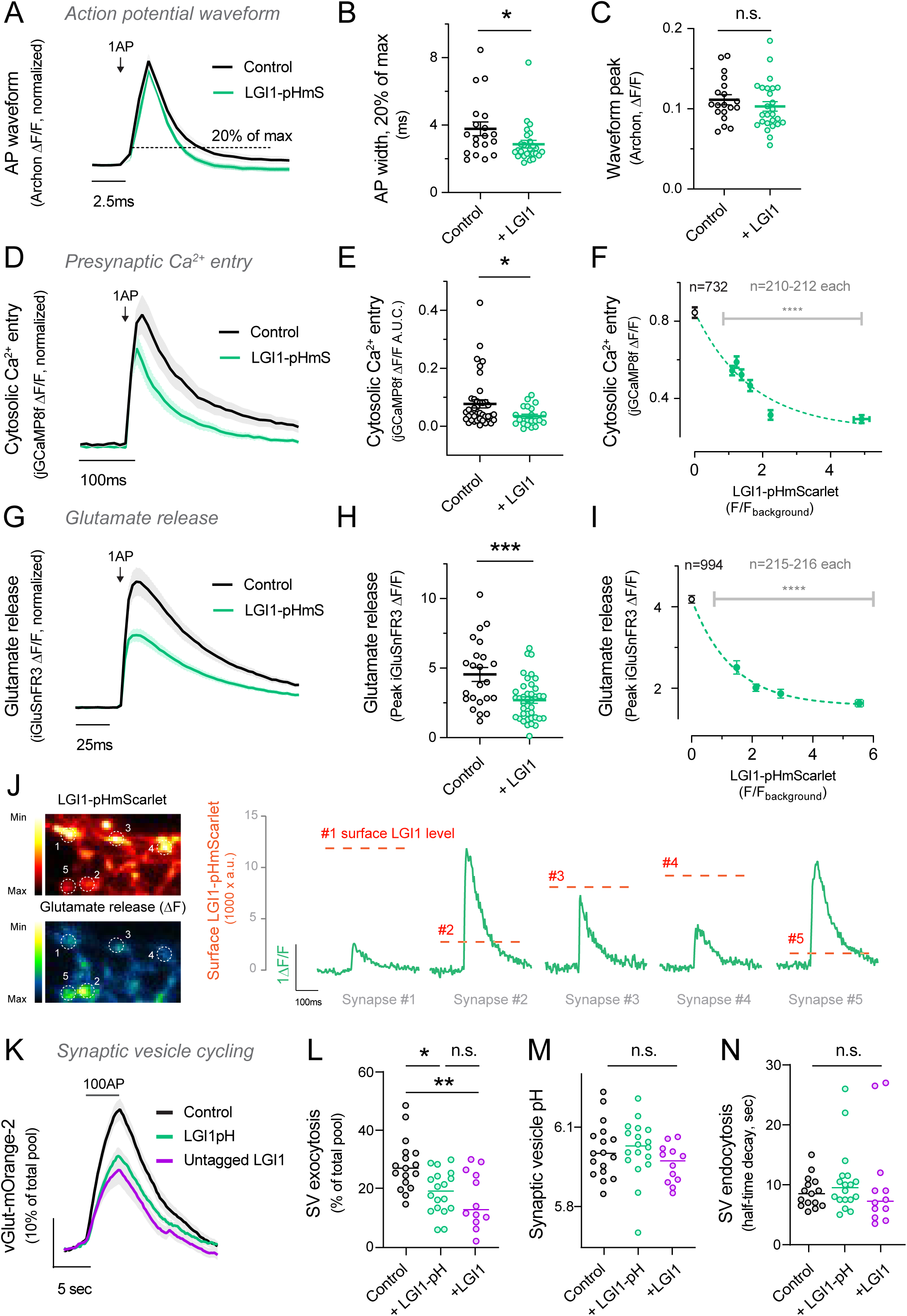
LGI1 abundance at the synaptic cleft controls the presynaptic action potential waveform, calcium entry, synaptic vesicle cycling and glutamate release. (**A**) Presynaptic action potential waveform measured using Archon in control versus LGI1-pHmS overexpressing neurons. **(B)** Comparison of the AP width by quantification of the full width at 0.2 of the peak response of each neuron. Data are mean ± SEM. Control n=19, 3.78 ± 0.41; LGI1 overexpression, n=26, 2.87 ± 0.23. **(C)** Quantification of peak response of the action potential. Data are mean ± SEM. Control n=19, 0.111 ± 0.006; LGI1 overexpression, n=26, 0.103 ± 0.006. Mann Whitney test. *p<0.05. **(D)** Average of cytosolic calcium entry to 1AP stimulus in control versus LGI1-pHmS overexpressing neurons. n=40 and n=21 neurons, respectively. Bars are SEM. **(E)** Quantification of D; n=40 and n=21 neurons. **(F)** LGI1-pHmScarlet expression and corresponding Ca^2+^ entry during 1AP stimulus was performed by grouping individual boutons according to their LGI1-pHmScarlet fluorescence and averaging their corresponding 1AP-evoked Ca^2+^ entry. Binning size was fixed at n=212 per group for equivalent sampling, except for controls not co-transfected with LGI1-pHmScarlet (n=732). Data are mean ± SEM. Group 0, n=732, LGI1-pHmScarlet 0.00 ± 0.00, jGCaMP8f 0.84 ± 0.03; group 1, n=210, LGI1-pHmScarlet 1.10 ± 0.003, jGCaMP8f 0.54 ± 0.02;; group 2, n=210, LGI1-pHmScarlet 1.23 ± 0.003, jGCaMP8f 0.59 ± 0.03; group 3, n=212, LGI1-pHmScarlet 1.379 ± 0.004, jGCaMP8f 0.523 ± 0.028; ; group 4, n=212, LGI1-pHmScarlet 1.63 ± 0.01, jGCaMP8f 0.46 ± 0.03; ; group 5, n=212, LGI1-pHmScarlet 2.24 ± 0.02, jGCaMP8f 0.31 ± 0.03; ; group 6, n=211, LGI1-pHmScarlet 4.92 ± 0.23, jGCaMP8f 0.29 ± 0.02. Dotted line represents the fitting of the data to a single exponential decay model. Dunn’s multiple comparisons test of the Group 0 vs. Group 1, showed no statistical difference (n.s.); Group 0 vs. Group 2 ****p<0.0001; Group 0 vs. Group 3 ****p<0.0001; Group 0 vs. Group 4 ****p<0.0001; Group 0 vs. Group 5 ****p<0.0001; Group 0 vs. Group 6 ****p<0.0001. **(G)** Average traces showing glutamate release in response to 1AP stimulus in control versus LGI1-pHmScarlet overexpressing neurons. n=23 and n=40 neurons, respectively. Bars are SEM. **(H)** Quantification of glutamate release to 1AP stimulus in control versus LGI1 overexpressing neurons. n=23 and n=40 neurons. **(I)** LGI1-pHmScarlet expression anticorrelation to presynaptic glutamate release in single boutons was performed as in F, using a binning size was fixed at n=216 per group for equivalent sampling, except for controls not co-transfected with LGI1-pHmScarlet (n=994). Data are mean ± SEM. Group 0, n=994, LGI1-pHmScarlet 0.00 ± 0.00, iGluSnFR3 4.18 ± 0.09; group 1, n=215, LGI1-pHmScarlet 1.49 ± 0.02, iGluSnFR3 2.51 ± 0.16; group 2, n=216, LGI1-pHmScarlet 2.13 ± 0.01, iGluSnFR3 2.02 ± 0.09; group 3, n=216, LGI1-pHmScarlet 2.94 ± 0.02, iGluSnFR3 1.87 ± 0.11; group 4, n=216, LGI1-pHmScarlet 5.54 ± 0.13, iGluSnFR3 1.63 ± 0.09. Dunn’s multiple comparisons test of the Group 0 vs. Group 1 ****p<0.0001; Group 0 vs. Group 2 ****p<0.0001; Group 0 vs. Group 3 ****p<0.0001; Group 0 vs. Group 4 ****p<0.0001. **(J)** Representative example image of LGI1-pHmScarlet baseline signal (top left) and glutamate release (ΔF) (bottom left) in different synaptic boutons of a single neuron (1-5). Glutamate release traces for each individual bouton from 1 to 5 (green) and the associated surface LGI1-pHmScarlet level (dotted line, scale on the left) exemplify the correlation observed in (I). **(K)** Average traces showing synaptic vesicle cycling during 100AP 10Hz in control (n = 17 neurons) versus LGI1-pH (n = 18 neurons) or untagged LGI1 (n = 12 neurons) overexpression. **(L)** Quantification of synaptic vesicle exocytosis peaks in response to 100AP fired at 10Hz in control versus neurons overexpressing LGI1-pH or untagged LGI1. Exocytosis is measured as percentage of SV pool mobilized by stimulation, where 100% is the signal obtained by applying NH_4_Cl pH 7.4 Data are mean ± SEM. Control n=17, 28.0 ± 2.2; LGI1-pH n=18, 18.7 ± 1.7; LGI1 n=12, 16.2 ± 2.8. Dunn’s multiple comparisons test of the peaks of control vs. LGI1-pH *p<0.05; control vs. LGI1 **p<0.01; LGI1-pH vs. LGI1 n.s. **(M)** Synaptic vesicle pH estimated using vGlut-mOr2 in neurons expressing LGI1-pH or LGI1. Data are mean ± SEM in pH units, n=17 neurons, 6.01 ± 0.03 pH for control; n=18 neurons, 6.02 ± 0.03 for LGI1-pH; n=12 neurons, 5.96 ± 0.02 for LGI1. Dunn’s multiple comparisons test of control vs. LGI1-pH n.s.; control vs. LGI1 n.s.; LGI1-pH vs. LGI1 n.s. **(N)** Quantification of endocytosis rates after 100AP 10Hz stimulus, measured as the time to reach half the amplitude of the peak response (t half). Data are mean ± SEM. Control n=17, 9.35 ± 0.76; LGI1-pH n=18, 10.56 ± 1.28; LGI1 n=12, 10.17 ± 2.34. Dunn’s multiple comparisons test of the peaks of control vs. LGI1-pH n.s.; control vs. LGI1 n.s.; LGI1-pH vs. LGI1 n.s.

In our initial experiments we observed that synapses belonging to the same axon presented different levels of surface LGI1 (Figure 1E; Figure 4E; Supp. figure S1A) and thus we reasoned that if LGI1 is a key control mechanism of synaptic function, individual presynaptic strength in single nerve terminals could be correlatively modulated by local surface LGI1 abundance in synapses. As both jGCaMP8f and iGluSnFR3 allowed us to quantify single AP-driven responses in single synapses with sufficient signal-to-noise, we measured individual synapse surface levels of LGI1-pHmScarlet and the corresponding responses in Ca^2+^ entry and glutamate release to a single action potential. We first analyzed over 1300 individual boutons from 21 separate neurons expressing jGCaMP8f and LGI1-pHmScarlet and our data revealed a negative correlation in which higher expression of LGI1 was robustly correlated with lower AP-driven Ca^2+^ entry (Figure 6F). With a similar approach, we analyzed both LGI1-pHmScarlet fluorescence and single AP-driven glutamate release in individual boutons. We analyzed over 800 individual boutons from 40 separate neurons and we confirmed that presynaptic sites with higher expression of LGI1 at the surface released significantly less glutamate during single action potential firing (Figure 6F). Such modulation was easily identifiable in individual boutons from presynaptic arborizations of single neurons (Figure 6J), supporting the analysis shown in Figure 6F.

As LGI1 modulates action potential waveform, Ca^2+^ entry and glutamate release, we next explored the role of LGI1 in controlling synaptic vesicle cycling. We co-transfected red-shifted vGlut-mOrange-2^65^ with LGI1-pH and tested synaptic vesicle cycling during a train of 100AP triggered at 10Hz. As expected, we found that exocytosis was reduced by ∼30% when LGI1-pH was overexpressed (Figure 6K, L). We also used this paradigm to confirm that the presence of pHluorin in the LGI1-pH construct does not alter the function of LGI1. We thus tested the impact of overexpressing untagged LGI1 onto synaptic vesicle cycling and observed an impairment in exocytosis indistinguishable from that obtained using LGI1-pH, suggesting that LGI1 function is not impaired by tagging it with pHluorin. Similarly, we confirmed that the presence of LGI1-pH nor LGI1 does not alter synaptic vesicle properties, including synaptic vesicle pH (Figure 6M), endocytosis rates (Figure 6N) or synaptic vesicle pool size (Supp. Figure S9A). While overexpressing LGI1 could be also altering neurotransmitter loading into synaptic vesicles, the fact that we observe alterations in action potential waveform, Ca^2+^ entry and synaptic vesicle exocytosis is more easily explained by a modulation of presynaptic exocytosis than reduced neurotransmitter loading. Taken together, results in Figure 6 show that LGI1 surface abundance in individual boutons controls locally the action potential waveform and Ca^2+^ entry, serving as a major modulatory mechanism of glutamate release that controls differentially presynaptic strength in boutons belonging to the same axon.

### Patient-derived autoantibodies against LGI1 reduce its surface abundance and increase glutamate release

Autoantibodies against LGI1 cause a form of limbic encephalitis (LE) associated with cognitive decline and seizures^11–14^, although the mechanisms by which these antibodies cause neuronal dysfunction remains poorly understood. In several other autoimmune neurological disorders, autoantibodies bind to their target receptors and cause their internalization and subsequent loss of function^66,67^. We reasoned that autoantibodies against LGI1 could induce a reduction in surface abundance of LGI1 at synapses and leveraged the use of our novel tools to test this idea. We obtained plasma from patients suffering LE together with control plasma (see STAR methods), purified IgG antibodies and treated primary cultures expressing LGI1-pH for a week in the presence of purified IgG from both conditions (Figure 7A). Using our methods to quantify the fraction of LGI1 molecules present at the synaptic surface, we found that the presence of autoantibodies reduced LG1-pH at the synaptic surface by ∼50% when compared to neurons treated with control IgGs (Figure 7B, C). This result shows for the first time a direct measurement supporting that anti-LGI1 antibodies reduce the presence of LGI1 function at the synaptic surface, thus impairing its function. We confirmed that the total amount of LGI1-pH remained unaltered (Figure 7D) in these conditions. We hypothesized that, contrary to our overexpression experiments (Figure 6), reducing the presence of LGI1 at the synaptic surface should result in increased glutamate release^20^. To test this, we treated primary neurons expressing iGluSnFR3 with control or LE IgGs as before and measured AP-induced glutamate release. We found that impairing the function of endogenous LGI1 resulted in significantly higher glutamate release, causing a ∼45% increase (Figure 7E, F). The presence of LE IgGs did not modulate glutamate clearance rates at the presynapse (Supp. Figure S9B), suggesting an effect solely on glutamate release. Taken together, these experiments show that modulating the presence of endogenous LGI1 at the synaptic surface results in a corresponding change in glutamate release and provide the first experimental evidence suggesting that autoantibodies against LGI1 exert their pathological effect by driving a reduction of surface LGI1 in neurons.

**Figure 7.**
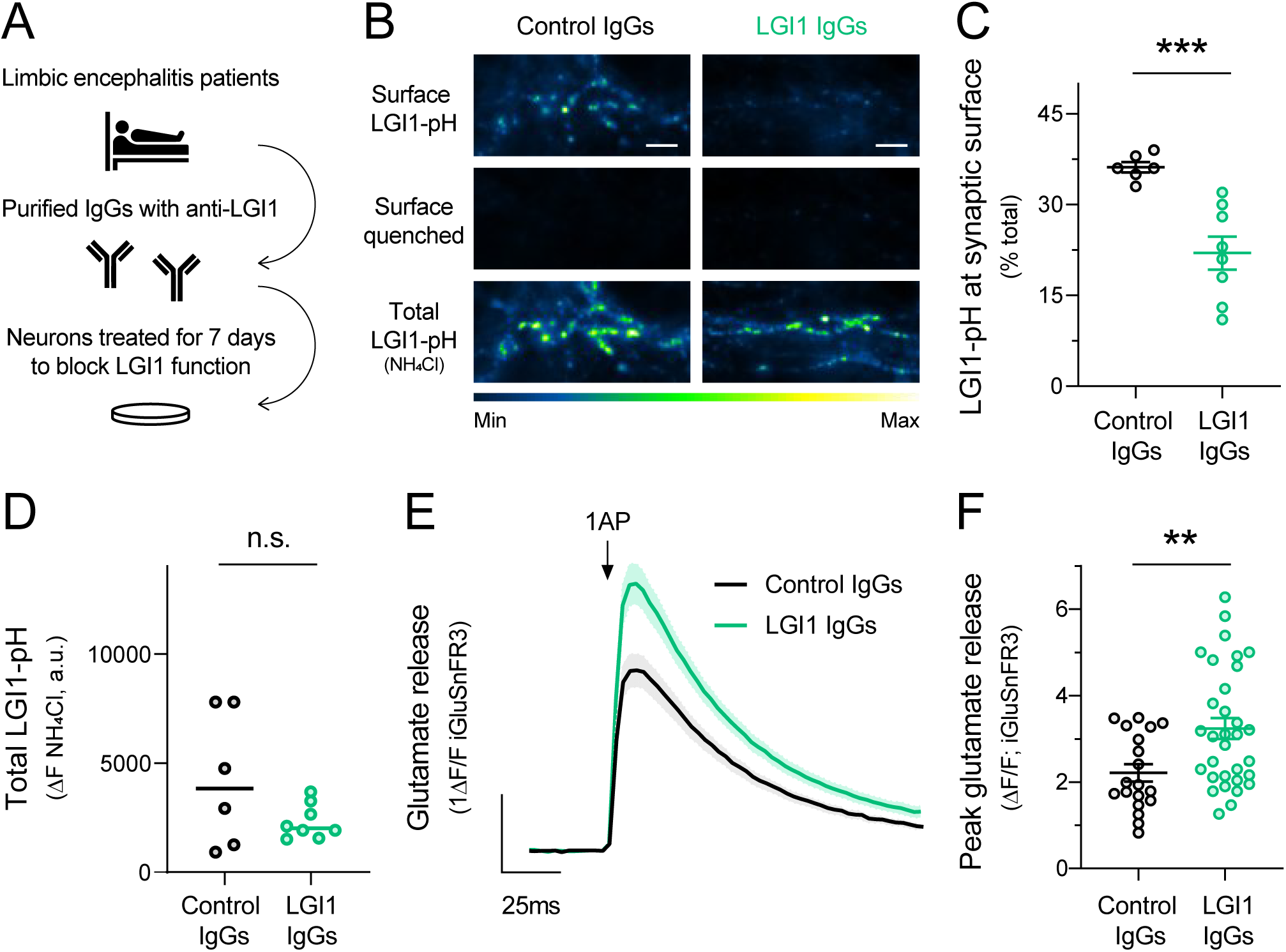
Anti-LGI1 antibodies reduce LGI1 surface fraction and cause increased glutamate release. (**A**) Graphic scheme of the protocol followed to treat primary neurons with anti-LGI1 auto-antibodies obtained from plasma of limbic encephalitis patients. **(B)** Example images of LGI1-pH expression levels at the surface (top panel), quenched surface LGI1-pH (middle panel, quenched by application of MES Tyrode’s at pH 5.5) and total LGI1-pH (bottom panel, revealed by NH_4_Cl pH 7.4) in neurons treated with control IgGs or anti-LGI1 patient-derived IgGs. Scale bar 7.6μm. **(C)** Quantification of LGI1 abundance in synaptic surface of neurons expressing LGI1-pH and treated with either IgGs from control or anti-LGI1 IgGs from limbic encephalitis patients. n=6 and n=8 neurons, respectively; Data are mean ± SEM. Control IgGs 36.2 ± 0.9; LGI1 IgGs 22.0. ± 2.7. t-test, ***p<0.001. **(D)** Total LGI1-pH in neurons incubated with control IgGs versus LGI1 IgGs, measured as peak response in NH4Cl pH 7.4. Data are mean ± SEM. Control IgGs n=6, 4240 ± 1254; LGI1 IgGs n=8, 2325 ± 282. t-test, n.s. **(E)** Average traces showing glutamate release in response to 1AP stimulus in neurons treated with IgGs from control or anti-LGI1 IgGs from limbic encephalitis patients. n=19 and n=32 neurons, respectively. Bars are SEM. **(F)** Quantification of individual peak responses shown in E. Data are mean ± SEM. Control IgGs n=19; 2.21 ± 0.21; anti-LGI1 n=32; IgGs 3.24. ± 0.24. t-test, **p<0.01.

## Discussion

A decade of accumulating evidence using *in vivo* experimental approaches has demonstrated that reductions in LGI1 drive synaptic dysfunction, causing seizures and premature death in rodents. However, how the presence of LGI1 at the synaptic surface is controlled has remained poorly understood. Leveraging the development of novel optical tools to monitor the behavior of LGI1 in single firing synapses, we found that neuronal activity is a robust controller of LGI1 abundance at the synaptic cleft. The history of activity of a neuron is essential for developing appropriate connectivity^68^, which requires the formation of strong trans-synaptic interactions to facilitate the function of established synapses^9,69,70^. We find that both chronic and acute neuronal activity stably boost LGI1-mediated trans-synaptic bridges and their abundance at the cleft of individual terminals correlatively attunes presynaptic function. Given that optimal synaptic transmission relies heavily on the sub-synaptic molecular architecture^9^, our results suggest that neuronal activity may acutely remodel trans-synaptic molecular landscape to adapt the strength of the connection. Indeed, activity remodels synaptic geometry^71^, driving structural changes that could be the consequence of activity-driven adjustments of trans-synaptic molecular networks.

LGI1 is broadly considered as a secreted protein, as when overexpressed in heterologous cells it can easily be found in their media^27–33^. Applying purified LGI1 obtained from heterologous cell media impairs Kv1 function and reduces the intrinsic excitability of CA3 neurons in slices^19^, in agreement with the idea that soluble LGI1 can bind available ADAM22/23 receptors^14,18^. However, while LGI1 can also be detected in the media of cultured neurons and organotypic slices^25,45^, this occurs to much less extent than in heterologous cells^25^. In agreement with the latter observation, our data shows that the majority of LGI1 molecules exposed during firing do not behave as secreted proteins, but are trafficked to, and retrieved from, the synaptic surface. We propose this is possible through a sustained interaction with ADAM23, which also is exposed to the synaptic surface with similar dynamics to LGI1 during activity. Contrarily, we can detect LGI1 secretion if its interaction with ADAM23 is impaired by the LGI1 mutation Y433A^16^. Similarly to what occurs in heterologous cells, which do not express ADAM23^27–33^, we propose that LGI1^Y433A^ cannot be fully retained by ADAM23 in synapses and thus behaves partially as a secreted protein (Figure 3F, G). Taken together, our data indicate that while LGI1 is indeed a secreted protein, and it can behave as such, it is unlikely that it is being significantly secreted during its translocation at synapses. This unexpected result challenges the current notion on how LGI1, and possibly other secreted synaptic proteins, are translocated to the synaptic cleft and exert their function.

Mechanisms preventing secretion of LGI1 optimize the metabolic efficiency of its translocation, as the energy spent in its synthesis is not lost if LGI1 molecules that are not successful in forming a trans-synaptic bridge can be recovered back to the terminal through endocytosis. Indeed, evolutionary pressure has optimized neuronal function to favor metabolic efficiency^72–75^, and our results highlight yet another mechanism that ameliorates the presynaptic energetic burden associated with neuronal activity^37,76,77^. Moreover, given that on average an LGI1 vesicle will contain 6 LGI1 molecules (Supp. Figure S4C) but only a fraction stabilizes at the synaptic cleft after activity (Figure 4A, B), recovering the non-stabilized molecules back to the presynapse should enable subsequent rounds of LGI1 translocation without requiring novel LGI1 synthesis. We find a relatively high probability of multivesicular exocytosis during trains of action potentials at 50Hz during 20 seconds (Supp. Figure S2A), which correspondingly translocate on average ∼30% of the total pool of LGI1 molecules to the synaptic surface (Figure 2G). We hypothesize that if that amount of LGI1 was not endocytosed but secreted and lost after translocation, LGI1 protein levels would almost disappear in about ten rounds of firing at those frequencies (see simulation in Supp. Figure S9C). The fact that no LGI1-pH loss is observed experimentally during action potential firing (Figure 3D, E) suggests that synapses recover translocated LGI1 molecules by endocytosis, contributing to preserving presynaptic LGI1 pools in the short term. This would liberate synapses from continuously relying on strong *de novo* LGI1 synthesis and transport from neuronal somas, which are typically located millimeters to centimeters away from the synaptic site.

Taken together, our data indicates that LGI1 does not behave as a canonical secreted protein. This is supported by A) the fact that dynamics of LGI1 translocation differ significantly from the dynamics of a canonical secreted protein such as Neuropeptide-Y (Figure 3A-C) and B) the fact that strong stimulation paradigms translocate large amounts of LGI1 to the surface (Figure 2G) without incurring in LGI1 loss at the synapse (Figure 3D, E). Our results, however, are not incompatible with the fact that LGI1 is found secreted in the media of cultured primary neurons^25^ or organotypic slices^45^. First, our optical assays reveal a small proportion of LGI1-pH molecules being secreted or dissociated from synaptic receptors (Supp. Figure S3C, D), which over time could lead to the detectable presence of LGI1 in the neuronal medium. Second, recent work has shown that 71% of the neuronal secretome is originated by proteolytic shedding of membrane proteins rather than vesicular secretion^45^, indicating that the presence of a protein in the extracellular medium does not necessarily imply it was secreted. Moreover, this work demonstrated that the ectodomain of ADAM22 (ECD), which is necessary for LGI1 binding at the postsynapse, is constitutively cleaved, suggesting that such shredding should release of both ECD-ADAM22 and LGI1 that was bound to it into the media^45^.

Given the clinical importance of LGI1, there is a critical need to develop novel strategies to establish the pathogenicity of newly identified LGI1 genetic variants in the clinic. As current assays only distinguish qualitatively between secretion-competent and -incompetent mutants, deciphering whether a secretion-competent mutant is pathogenic requires laborious experimentation, including co-immunoprecipitation and binding assays to ADAM22/23 receptors^16,32^. We provide here a simple quantitative toolkit to robustly test several possible aspects of dysfunction in LGI1 by combining the use of primary neurons and field stimulation. We demonstrate that this approach reveals defects that were previously undetectable for epilepsy-associated mutants: 1) LGI1^T380A^, despite defined previously as secretion-incompetent^27^, translocates partially to the synaptic surface and is partially retained in axonal ER and 2) LGI1^S473L^ and LGI1^R474Q^, while translocate efficiently, they cannot get stably increased in synapses after neuronal activity. Similarly, our tools helped to dissect quantitatively how autoantibodies against LGI1, which cause limbic encephalitis (LE) and seizures^11,13^, can affect the abundance of LGI1 at the synaptic surface and impair synaptic function. We found that autoantibodies drive an internalization of LGI1 from the synapse surface, resulting in an increased capacity of glutamate release. These results provide the first direct evidence, to our knowledge, indicating that autoantibodies against LGI1 cause pathology by removing LGI1 molecules from the synaptic surface. This phenotype aligns with results observed in several other autoimmune diseases of the nervous system^66,67^ and supports the idea that autoantibodies against LGI1 induce the internalization of the protein at synapses, leading to increased glutamate release that could change the excitation-inhibition balance and cause seizures. Thus, the quantitative nature of this novel methodology provides the field with an improved assay for better understanding the molecular dysfunction of LGI1 and a novel approach for dissecting the pathogenicity of new mutations identified in the clinic.

While it is known that LGI1 dysfunction leads to disease through increased brain excitation, controversy remains on the cellular origins of this imbalance^4,23^. Here we find that the abundance of surface LGI1 in single synapses strongly influences the presynaptic action potential waveform, presynaptic Ca^2+^ handling, synaptic vesicle exocytosis and glutamate release (Figure 6). Remarkably, we find that such control is synapse-specific. Different presynaptic sites belonging to the same axon present different levels of surface LGI1, which strongly correlate with their variable presynaptic strength (Figure 6J). While in these experiments we modulated LGI1 abundance through overexpression, we found the opposite effect when we reduced the presence of endogenous LGI1 at the synaptic surface using anti-LGI1 antibodies: a reduced the amount of LGI1 at synapses (Figure 7B, C) correspondingly drove an increase in glutamate release (Figure 7E, F). This suggests that neurons could modulate surface LGI1 levels to functionally adapt glutamatergic transmission in healthy states, but if pathological states alter LGI1 localization at the synaptic surface, those synapses will present excessive glutamatergic transmission, causing circuit imbalance and seizures.

Computational complexity in the brain is thought to benefit from the diversity in presynaptic strength found in diverse neuronal connections^78,79^ and in different presynaptic sites belonging to the same axon^80–82^. Principal neurons like pyramidal cells of the hippocampus can present up to 15.000 synaptic contact sites^83^, and the ability to individually tune their strength increases the complexity of information that can be transmitted^84^. However, the molecular underpinnings enabling such heterogeneity in presynaptic strength, and how such variability is readjusted during different functional states, remain poorly understood. Our work identifies LGI1 as a novel regulator of presynaptic strength variability and, surprisingly, this modulatory role appears not to be static but adjustable in short timescales. This enables LGI1 to act as an integrator of synaptic function that correlatively attunes synaptic strength to match the history of activity to neurotransmission.

The fact that both LGI1 and its presynaptic receptor ADAM23 can acutely change their abundance at the synaptic cleft during activity suggests that the molecular landscape of the synaptic cleft is plastic in faster timescales than initially thought^85^. As these trans-synaptic proteins are not present in typical synaptic vesicles (Figure 2), our results open up the possibility that the surface abundance of other presynaptic proteins may be regulated by the exocytosis of these alternative vesicles. For example, the presynaptic translocation of a glucose transporter, GLUT4, has been shown to occur from presynaptic vesicles that are not SVs and whose luminal pH is significantly less acidic^37^, as we find for LGI1 (Figure 2A). Thus, future studies dissecting the proteome of LGI1 vesicles surely will provide new insights into defining the dynamic nature of synaptic cleft proteome and the extent to which the surface abundance of certain molecules may modulate synaptic physiology. Taken together, our results open new avenues of research that will define how activity can remodel the trans-synaptic molecular landscape of firing synapses in fast timescales, defining novel molecular mechanisms controlling neurotransmission.

## STAR Methods

### Resource availability Lead contact

Further information and requests for resources and reagents may be directed to and will be fulfilled by the lead contact, Dr. Jaime de Juan-Sanz.

### Material availability

Novel optical tools generated in this work are available at Addgene.org, as indicated in the Key Resources Table.

| REAGENT or RESOURCE | SOURCE | IDENTIFIER |
| --- | --- | --- |
| Antibodies |  |  |
| Rabbit anti-Lgi1/EPT antibody | Abcam | ab30868 |
| Mouse anti-GRIA1 antibody | UC Davis/NIH NeuroMab Facility | 75-327 |
| Goat Anti-Rabbit IgG (H + L)-HRP Conjugate | BioRad | 1706515 |
| Goat Anti-Mouse IgG (H + L)-HRP Conjugate | BioRad | 1706516 |
| Chemicals, peptides, and recombinant proteins |  |  |
| Bafilomycin A<sub>1</sub> | Cayman Chemical Company | Item #11038 |
| EGTA-AM | Thermo Fisher Scientific | E1219 |
| Tetrodotoxin citrate (TTX) | Tocris | Cat #1069 |
| Biotin-XX Tyramide Reagent (BxxP) | ApexBio Technology | Cat #A8012 |
| Tetanus toxin from Clostridium tetani | Sigma | T3194 |
| Recombinant Enhanced GFP protein (His tag) | Abcam | ab134853 |
| Critical commercial assays |  |  |
| Q5® Site-Directed Mutagenesis Kit | New England Biolabs | E0552 |
| Pierce™ High Sensitivity Streptavidin-HRP | Thermo Fisher Scientific | 21130 |
| NEBuilder® HiFi DNA Gibson Assembly Master Mix | New England Biolabs | E2621 |
| Experimental models: Organisms/strains |  |  |
| Sprague-Dawley Rat | Charles River | Strain: Crl:CD(SD) |
| Recombinant DNA |  |  |
| CamKII-LGI1-pHluorin | This paper | Addgene Plasmid #185537 |
| CamKII-LGI1-S473L-pHluorin | This paper | Addgene Plasmid #185540 |
| CamKII-LGI1-Y433A-pHluorin | This paper | Addgene Plasmid #185541 |
| CamKII-LGI1-R474Q-pHluorin | This paper | Addgene Plasmid #185542 |
| CamKII-LGI1-T380A-pHluorin | This paper | Addgene Plasmid #185543 |
| CamKII-LGI1-E383A-pHluorin | This paper | Addgene Plasmid #185544 |
| CamKII-LGI1-C200R-pHluorin | This paper | Addgene Plasmid #185545 |
| CamKII-LGI1 (no tags) | This paper | N/A |
| vGlut-pH | Gift from Timothy A. Ryan | N/A |
| vGlut-mOrange-2 | Hoppa et al., 2012<sup>80</sup> | N/A |
| iGluSnFR3 | Addgene | Addgene Plasmid #178330 |
| CMV-mRuby-synapsin | Addgene | Addgene Plasmid #187896 |
| pKanCMV-mRuby3-18aa-actin | Addgene | Addgene plasmid #74255 |
| pLKO-synapsin-BFP | Addgene | Addgene Plasmid #191567 |
| pLKO-empty-mTagBFP | Addgene | Addgene Plasmid #191566 |
| CamKII-ADAM23-pHluorin | This paper | Addgene Plasmid #185539 |
| pGP-AAV-syn-jGCaMP8f-WPRE | Addgene | Addgene Plasmid #162376 |
| CamKII-LGI1-pHmScarlet | This paper | Addgene Plasmid #185538 |
| CamKII-LGI1-R474Q-pHmScarlet | This paper | N/A |
| CamKII-LGI1-T380A-pHmScarlet | This paper | N/A |
| CamKII-LGI1-E383A-pHmScarlet | This paper | N/A |
| pCAGGS-SNAP25ΔC-IRES-mCherry | Ibata et al., 2019 | N/A |
| pCAGGS-SNAP29ΔC-IRES-mCherry | Ibata et al., 2019 | N/A |
| pCAGGS-Stx1AΔC-IRES-mCherry | Ibata et al., 2019 | N/A |
| pCAGGS-Stx4ΔC-IRES-mCherry | Ibata et al., 2019 | N/A |
| CMV-IgK-pHluorin-TM-mRuby | This paper | Addgene Plasmid #185546 |
| ER-pHluorin | de Juan-Sanz et al., 2017<sup>36</sup> | N/A |
| FSW-HRP-V5-LRRTM2 | Addgene | Addgene Plasmid #82537 |
| FSW-HRP-V5-LRRTM1 | Addgene | Addgene Plasmid #82536 |
| pAAV-CaMKII-Archon1-EGFP | Addgene | Addgene Plasmid #108417 |
| hSyn-NPY-pHluorin | Gift from Ruud F Toonen<sup>86</sup> | N/A |
| Software and algorithms |  |  |
| ImageJ | National Institute of Health | https://imagej.nih.gov/ij/ |
| GraphPad Prism 8 | GraphPad Software | https://www.graphpad.com/e/prism/ |
| R | R-project | https://www.r-project.org |
| RStudio | RStudio | https://www.rstudio.com |

### Experimental model and subject details

#### Animals

Wild-type rats were of the Sprague-Dawley strain Crl:CD(SD), which are bred by Charles River Laboratories worldwide following the international genetic standard protocol (IGS). All experiments were carried out in strict accordance with the guidelines of the European Directive 2010/63/EU and the French Decree n° 2013-118 concerning the protection of animals used for scientific purposes.

#### Primary co-culture of postnatal neuronal and astrocytes

All experiments were performed in primary co-cultures of hippocampal neuronal and astrocytes, except for proximity biotinylation experiments. P0 to P2 rats of mixed-gender were dissected to isolate hippocampal CA1-CA3 neurons. Neurons were plated on poly-ornithine-coated coverslips, transfected 5-7 days after plating, and imaged 14-21 days after plating. Transfection was performed using the calcium phosphate method, as previously described^34^, using equal amounts of DNA when co-transfecting two different plasmids. This method yields a very sparse transfection in primary neuronal cultures, with a 0.5-1% efficiency that allows to study fluorescence changes in presynaptic arborizations of single neurons during field stimulation. Neurons were maintained in culture media composed MEM (Thermo Fisher Scientific), of 20 mM Glucose, 0.1 mg/mL transferrin (Sigma), 1% Glutamax (Thermo Fisher Scientific), 24 μg/mL insulin (Sigma-Aldrich), 5% FBS (Thermo Fisher Scientific), 2% N-21 (Bio-techne) and 4μM cytosine β-d-arabinofuranoside (Milliopore). Cultures were incubated at 37°C in a 95% air/5% CO_2_ humidified incubator for 14– 21 days prior to use.

#### Primary culture of embryonic cortical neurons for synaptic cleft biotinylation

Pregnant wild-type Sprague Dawley Crl:CD(SD) rats were purchased from Charles River Laboratories. After euthanasia using CO_2_ (carried out by the animal facility from Paris Brain Institute), embryos were sacrificed at embryonic day 18 (E18). Dissected rat embryo cortical tissue was digested with papain (Worthington Biochemical, LK003178) and DNase I (Roche, 10104159001) and 3.5 million cortical neurons were plated per 10 cm dish, which were previously coated through a pre-incubation overnight at 37 °C with 0.1 mg/ml poly-D-lysine (Sigma, P2636). Neurons were cultured at 37°C under 5% CO_2_ in plating medium composed by a 1:1 volume ratio of growth medium A and growth medium B. Growth medium A is composed of: MEM (Sigma) with L-glutamine (Sigma) supplemented with 10% (v/v) fetal bovine serum (GIBCO) and 2% (v/v) N21 (Life Technologies). Growth medium B is composed of: Neurobasal medium (Life Technologies) supplemented with 2% (v/v) N-21 (Bio-techne) and 1% (v/v) GlutaMAX (Sigma). At 5 days in vitro, half of the culture medium was replaced with fresh BrainPhysTM Neuronal Medium supplemented with 2% (v/v) SM1 (STEMCELLTM Technologies, 05792) and 12.5 mM D-(+)-Glucose (Sigma, G8270) in addition to 10 µM 5’-fluoro-2’-deoxyuridine (FUDR, Fisher Scientific, 10144760), an antimitotic drug less toxic than the commonly used Cytosine Arabinofuranoside (Ara-C). While Ara-C does not impact the health of astrocyte-neuron co-cultures used for imaging, we found that it appeared to be excessively toxic for pure neuronal cultures. In contrast, FUDR provided pure neuronal cultures that could be maintained without apparent toxicity for more than three weeks and thus became our preferred antimitotic drug for pure neuronal cultures. This medium replacement was repeated every 4-5 days until the day of the experiment.

#### Method details

*Replication:* Not applicable

*Strategy for randomization and/or stratification:* Not applicable

*Blinding at any stage of the study:* Analysis was not blinded to genotype *Sample-size estimation and statistical method of computation:* Not applicable

#### Inclusion and exclusion criteria of any data or subjects

Glutamate release and AP-driven presynaptic Ca^2+^ signals in response to electrical activity (ΔF) were normalized to the resting fluorescence (F_0_). To avoid overestimating ΔF/F_0_ in responding neurons with low F_0_ values, we set an arbitrary threshold such that F_0_/background > 1.25 to be included for further analysis^36^. The rationale for using such a threshold is as follows: a neuron expressing a fluorescent sensor must present enough signal at baseline fluorescence (F_0_) over background fluorescence (F_background_) for accurate quantification of ΔF/F_0_ responses. If F_0_ and F_background_ are too similar, as the difference between F_0_ and F_background_ approaches 0, the quotient of ΔF divided by (F_0_-F_background_) approaches infinity and thus ΔF/F_0_ estimates are overestimated in a non-linear fashion. This can be curbed partially by excluding neurons that are not at least 25% brighter than the background. This threshold did not exclude any iGluSnFR3 or jGCaMP8f responses obtained from single presynaptic arborizations. However, when applied to single-bouton responses, 51 out 915 iGluSnFR3 responses were excluded (5.5%), while no response was excluded for jGCaMP8f experiments. For the experiments quantifying LGI1-pHluorin and ADAM23-pHluorin changes during prolonged stimulation (1000AP 50Hz, 3000AP 50Hz) we occasionally observed a global decrease in fluorescence of the entire field of view, including the background. Thus, we set a threshold to exclude experiments without stable background conditions. Background regions were measured over time and experiments that experimented a change in background higher than 30% were excluded. Thus, for 1000AP 50Hz stimulation recordings, 3 out of 21 neurons were excluded and for 3000AP 50Hz stimulation recordings, 2 neurons out of 10 were excluded. For experiments in which we analyzed single bouton LGI1-pH, NPY-pH and ADAM23-pH exocytosis responses, we analyzed responses whose change in ΔF/F was at least 6 times the standard deviation of the baseline before exocytosis.

#### Gene constructs

LGI1-pHluorin was designed to express wild type rat LGI1 followed by a short flexible linker (SGSTSGGSGGTG) and pHluorin. This construct was optimized *in silico* for rat protein expression, synthesized *in vitro* (Invitrogen GeneArt Gene Synthesis) and cloned into the BamHI and EcoRI sites of the CaMKII promoter vector for exclusive expression in excitatory neurons^38^. CamKII vector was a gift from Edward Boyden (Addgene plasmid #22217). LGI1 E383A mutant was generated on this vector using site-directed mutagenesis (Q5® Site-Directed Mutagenesis Kit, New England Biolabs E0552) following manufacturer instructions using the following primers: Primer sequence 5’ - 3’ Forward: ACCGACGTCGCGTACCTGGAA, Primer sequence 5’ - 3’ Reverse: GTCTCTGTACCAGGCGTG. LGI1 mutants C200R, R474Q, S473L, Y433A in this vector were purchased at Eurofins. LGI1-pHmScarlet was created by replacing pHluorin with pHmScarlet using Gibson Assembly (NEBuilder® HiFi DNA Assembly Master Mix, New England Biolabs). Insert containing pHmScarlet was obtained by PCR amplification from VAMP2-pHmScarlet^55^, a gift from Pingyong Xu (Addgene plasmid # 166890). A plasmid to express mRuby-synapsin was generated by removing GFP from GFP-synapsin^87^ using restriction sites AgeI and BglII, and substituting it in frame with mRuby obtained from pKanCMV-mRuby3-18aa-actin, which was a gift from Michael Lin (Addgene plasmid # 74255). This plasmid was cloned in the laboratory of Timothy A. Ryan (Addgene plasmid # 187896). ADAM23-pHluorin was designed to express the signal peptide of wild type rat ADAM23 (first 56 amino acids) followed by pHluorin, a short flexible linker (SGSTSGGSGGTG) and the rest of ADAM23 (amino acids 56-828). This construct was optimized *in silico* for rat protein expression, synthesized *in vitro* (Invitrogen GeneArt Gene Synthesis) and cloned into the BamHI and EcoRI sites of the CaMKII promoter vector, which was a gift from Edward Boyden (Addgene plasmid #22217). These constructs have been deposited in addgene, where exact maps can be found easily.

#### Image analysis and statistics

We cultured and transfected primary hippocampal neurons in 30-40 independent coverslips per week. Each experiment was replicated in several independent imaging sessions at different days, as indicated in Supp. Table ST1. Image analysis was performed with the ImageJ plugin Time Series Analyzer V3 where typically 150-250 regions of interest (ROIs) corresponding to synaptic boutons or 10-150 ROIs for responding boutons were selected and the fluorescence was measured over time. Statistical analysis was performed with GraphPad Prism v8 for Windows. Statistic tests are indicated in the figure legends and in Supp. Table ST1. Mann–Whitney U test was used to determine the significance of the difference between two unpaired conditions without assuming normal distributions. Kruskal-Wallis test was used to determine whether there are any statistically significant differences between the medians of three or more independent groups. If a significant result from the Kruskal-Wallis test was obtained we used Dunn’s multiple comparison test to identify which specific groups differ from each other. Wilcoxon matched-pairs test was performed for our two paired data sets without assuming normal distributions. If datasets to be compared followed a normal distribution, which was evaluated by performing an Anderson-Darling normality test, we used Student’s t-test to compare two populations and one-way ANOVA with Dunnett’s multiple comparisons test for multiple populations. Throughout the text p<0.05 was considered significantly different and denoted with a single asterisk, whereas p<0.01, p<0.001 and p<0.0001 are denoted with two, three, and four asterisks, respectively. In experiments shown in Figures 3B, 4B and 5C we analyze paired comparisons of independent experiments to dissect whether individually each condition presents a significant change, instead of comparing the quantitative extent to which delta changes are different numerically. The rationale for this is that absolute numerical comparisons can be misleading if one needs to test relative changes in separate conditions that do not share the same underlying physiology, as is the case of different types of vesicles (Figure. 4), or different mutants (Figure 3, Figure 5). Throughout the text, when showing violin plots, quartiles are indicated by small dotted lines while the median is represented by a bold dotted line. When data are averaged, error shown represents SEM unless otherwise noted.

#### Live imaging of neurons

Primary hippocampal neurons were transfected using calcium phosphate at DIV7 as previously described^34^ and were imaged from DIV14 to DIV21. Imaging experiments were performed using a custom-built laser illuminated epifluorescence microscope (Zeiss Axio Observer 3) coupled to an Andor iXon Ultra camera (model #DU-897U-CSO-#BV), whose chip temperature is cooled down to -95°C using the Oasis™ UC160 Cooling System to reduce noise in the measurements. Illumination using fiber-coupled lasers of wavelengths 488 (Coherent OBIS 488nm LX 30mW) and 561 (Coherent OBIS 561nm LS 80mW) was combined through using the Coherent Galaxy Beam Combiner, and laser illumination was controlled using a custom Arduino-based circuit coupling imaging and illumination. Neuron-astrocyte co-cultures were grown in coverslips (D=0.17mm, Warner instruments), mounted on a RC-21BRFS (Warner Instruments) imaging chamber for field stimulation and imaged through a 40x Zeiss oil objective Plan-Neofluar with an NA of 1.30 (WD=0.21mm). Unless otherwise noted, imaging frequency was 2Hz. Different imaging frequencies are used in certain experiments: 100Hz for single action potential cytosolic jGCaMP8f, 350Hz for single action potential iGluSnFR3 and 2000Hz for single action potential waveform using Archon1. For voltage imaging, light was collected through the objective using a cropped sensor mode (10 MHz readout, 500 ns pixel shift speed) to achieve 2 kHz frame rate imaging (exposure time of 485 μs) using an intermediate image plane mask (Optomask; Cairn Research) to prevent light exposure of non relevant pixels as described previously^62^. To obtain sufficient signal to noise, each neuron AP waveform data comes from the average of 50 independent measurements taken at 5Hz. Temperature of all experiments was clamped at 36.5 °C, except for voltage imaging, where it was clamped at 34 °C. Temperature was kept constant by heating the stimulation chamber through a heated platform (PH-2, warner instruments) together with the use of an in-line solution heater (SHM-6, warner instruments), through which solutions were flowed at 0.35ml/min. Temperature was kept constant using a feedback loop temperature controller (TC-344C, warner instruments).

Imaging was performed in continuously flowing Tyrode’s solution containing (in mM) 119 NaCl, 2.5 KCl, 1.2 CaCl_2_, 2.8 MgCl_2_, 20 glucose, 10 mM 6-cyano-7-nitroquinoxaline-2,3-dione (CNQX) and 50 mM D,L-2-amino-5-phospho-novaleric acid (AP5), buffered to pH 7.4 at 37°C using 25 mM HEPES. NH_4_Cl solution for pHluorin measurements had a similar composition as Tyrode’s buffer except it contained (in mM): 50 NH_4_Cl and 69 NaCl for a pH of 7.4 at 37°C. The solution for surface acid quenching of pHluorin is identical to the Tyrode’s solution but was buffered using MES instead of HEPES and was set to pH 5.5 (at 37°C). Cells were flowed with MES and NH_4_Cl at faster speeds of 0.8ml/min for a quick pH change.

For experiments in figure 2, bafilomycin (Cayman Chemical Company) was diluted in Tyrode’s to a final concentration of 500nM and it was continuously flowed on neurons expressing either vGlut-pHluorin or LGI1-pHluorin for 1000 seconds, acquiring images every 0.5 seconds. In experiments in which neurons vGlut-pHluorin or LGI1-pHluorin transfected neurons were treated with EGTA-AM, we first acquired the response to stimulation, then neurons were incubated with 2mM EGTA-AM for 10 minutes, and then we acquired the response to stimulation in the same region. Chronic incubation with TTX (Tocris) was performed by adding 1 µM final concentration of TTX in the culture media 5-6 days prior imaging. Next, TTX was washed by flowing Tyrode’s solution during 10 minutes and optophysiological recordings using jGCaMP8f or iGluSnFR3 were performed.

#### pH, surface fraction and total pool measurements using pHluorin constructs

Intraluminal organelle pH in vGlut-, ADAM23- and LGI1-pHluorin-containing vesicles was calculated leveraging the known properties of pHluorin response to pH (pKa 7.1), as previously described^34^. The pH estimates obtained from LGI1-pH secretion-defective mutants C200R and E383A in axons was not reliable and thus it is not shown, as the signal-to-noise change in fluorescence during NH_4_Cl application was very low due to the low expression in the axon of these mutants. For measuring surface fraction of pH-tagged constructs, axons of neurons expressing vGlut-, ADAM23- and LGI1-pHluorin constructs were perfused briefly with an acidic solution at pH 5.5 buffered with MES (2-(N-morpholino)ethanesulfonic acid) for acid quench of pHluorin expressed at the neuronal surface followed by a NH_4_Cl solution at pH 7.4 for alkalization of vesicular pH, which reveals the total pool of pHluorin-tagged molecules. For these fast perfusions, flow rate was ∼0.8ml/min. Surface fraction of vGlut-pH, LGI1-pH and of LGI1 mutants-pH were determined before and after electrical stimulation using MES/NH_4_Cl measurements as previously described^34^. Cells were flowed sequentially with MES and NH_4_Cl solutions and then washed for 10min in Tyrode’s solution. Next, neurons were stimulated using field stimulation as indicated in the text (1000AP 50Hz or 3000AP 50Hz) and 5 minutes later surface fraction was measured again using MES/NH_4_Cl. To measure the total pool of LGI1 or LGI1 mutants before and after stimulation, a similar approach was used. Change in fluorescence by NH_4_Cl solution reveals the total pool present, and thus ΔF in the presence of NH_4_Cl pH 7.4 was acquired before and after electrical stimulation. All experiments were acquired with the same laser power and exposure times to be comparable.

#### Estimates of LGI1-pH molecules exocytosed per univesicular event using single EGFP imaging

Purified EGFP (Abcam) was diluted in PBS to a final concentration of 1 µg/ml and placed in a coverslip identical to those used for live neuron imaging. After droplets dried, we mounted 3 independent coverslips using ProLong (Thermo Fisher Scientific) and imaged 16 randomly selected fields at room temperature to quantify fluorescence corresponding to each EGFP dot identified, obtaining 1727 separate EGFP measurements. We fitted this population to a single gaussian distribution that presented an mean value of 535 a.u. (R-squared = 0.95), which we attribute to the fluorescence of a single EGFP molecule in our imaging conditions. We next imaged univesicular exocytosis of LGI1-pH during 200AP stimulation evoked at 50Hz using the same exact imaging conditions and analyzed 374 single-bouton exocytosis events obtained from 8 neurons. Changes in arbitrary units of fluorescence during exocytosis presented a median of 3015 a.u. (25% percentile = 1910; 75% percentile = 4594), which allows to conclude that approximately 6 LGI1-pH molecules are contained on average in a single LGI1 vesicle.

#### Identification of univesicular and multivesicular exocytosis events

The robustness of LGI1-pH responses allows studying single bouton events with a sufficient signal to noise that in a high fraction of the cases allows one to attribute fluorescence changes to univesicular or multivesicular exocytosis events. An exocytosis event is considered for our analyses if it fulfills the following criteria: 1) it presents a stable baseline during at least 3 seconds before the fluorescence increase and 2) it presents an increase of ΔF/F at least 6 times the standard deviation of that baseline before exocytosis.

To define whether events arise from the exocytosis of one or multiple vesicles, we first established the criteria for defining univesicular release. Univesicular events present a sharp increase In fluorescence in the sub second timescale, as exocytosis signals derived from pHluorins arise from the pH transition from ∼6.1 to 7.4 and the subsequent deprotonation of GFP, both of which occur in the millisecond timescale^88^. We thus consider an exocytosis event to be univesicular when the complete increase in fluorescence from baseline to maximum occurs in 1 second or less. This value is defined by the time resolution of our imaging frequency (2Hz). Supp. Figure S2B shows example responses.

Events were categorized as dual exocytosis events when during stimulation two complete increases in fluorescence from baseline to maximum occurred each in 1 second or less (see Supp. Figure S2C). Comparison of both events from the same presynaptic site revealed that on average fluorescence changes were identical in both the first and second events (Supp. Figure S4D, E). This indicates that our selection criteria for considering an exocytosis event as univesicular is accurate, as it would be very unlikely that an identical number of multiple vesicles are being exocytosed subsequently in the first and second events. Lastly, multivesicular events are those in which during stimulation, the fluorescence increases more than 6 times the value of the standard deviation of the baseline fluorescence but cannot be categorized as having only one or two exocytosis events following the criteria outlined above.

#### Identification of secretion-like responses in LGI1-pH translocation events

While LGI1-pH did not present dynamics resembling canonical secretion, in some cases translocation events presented mixed kinetics with partial secretion-like decreases in fluorescence (see Supp. Figure S3C). We identified these events in responding boutons after stimulation using the following criteria: 1) they presented a decrease in fluorescence of at least 3 times the standard deviation of the baseline before the event, 2) the decrease occurred in 1 second or less and 3) they presented a stable baseline for 2 seconds after the event (which was defined by excluding events whose fluorescence changed more than 10% after the sharp decrease). Possible events matching this criteria were initially identified manually to later quantify whether they fulfill the criteria stated above.

#### Delay to undergo exocytosis during electrical stimulation

To quantify the delay to respond during stimulation in individual boutons of neurons expressing vGlut-pH, LGI1-pH, NPY-pH or ADAM23-pH, we analyzed ΔF/F traces of individual boutons in each condition. We leveraged the sharp increase in fluorescence of responding boutons during stimulation to identify the time taken for responding since the stimulation began. To do so, we set a threshold for identifying a response: fluorescence of a responding bouton had to increase at least 6 times over the standard deviation of the baseline, which was calculated using the 20 time points before stimulation. To avoid including non-relevant fluctuations in fluorescence, boutons that did not maintain an increase in fluorescence of 6 times de standard deviation of the baseline during at least 1 second after the initial increase were excluded. Similarly, increases in fluorescence that were not larger than 6 times the standard deviation of the baseline were not part of this analysis. Individual asynchronous universicular exocytosis events of LGI1-pH, NPY-pH and ADAM23-pH, identified using the criterion explained above, were aligned to start rising at the same time to obtain average traces in figures 3A, 3B, 3L, 5D. We indicate sample sizes in each corresponding figure legend.

#### Quantification of endogenous surface LGI1 levels in the synaptic cleft by proximity biotinylation

Isolation of synaptic cleft proteins was performed as previously described in detail^52^, with small modifications. Two 10 cm dishes were plated with 3.5 million rat primary embryonic cortical neurons per experimental condition. At day 15 DIV, dishes were transduced with lentivirus expressing FSW-HRP-V5-LRRTM1 and FSW-HRP-V5-LRRTM2 constructs, using a total of 2 x 10^8^ VP/ml for transduction per 10 cm culture dish. After 4 days, live cell biotinylation experiments were performed at DIV19 as described below. Lentivirus were produced at the iVector facility at the Paris Brain Institute in BSL2 facilities transfecting HEK293T cells with constructs of interest together with 3^rd^ generation packaging, transfer and envelope plasmids, using the vesicular stomatitis virus G glycoprotein (VSVG) as envelope protein. Transient transfection of HEK293T was done using lipofectamine 2000 (Sigma-Aldrich) in a medium containing chloroquine (Sigma-Aldrich). The medium was replaced after 6 hours and the supernatant was collected after 36 hours. The supernatant was treated with DNAseI (Roche) and then ultracentrifugation was carried out at 22 000 rpm (rotor SW28, Beckman-Coulter) for 90 minutes. The resulting pellet was resuspended in 0.1M PBS, aliquoted and frozen at −80°C until use. Lentiviral productions presented titers ranging 9.9-19.7 x 10^8^ viral particles (VP) per μl, measured by Elisa using the p24 ZeptoMetrix kit (Sigma-Merck).

For evaluating the effect of activity in synaptic surface abundance of LGI1, chronic incubation with TTX (Tocris) was performed by adding 1 µM final concentration of TTX in the culture media 5 days prior to the experiment. Next, control and TTX-treated DIV19 neurons were exposed during 60 seconds to H_2_O_2_ and BxxP, an impermeant variant of biotin-phenol that contains a long and polar polyamide linker (ApexBio Technology), which allows selective biotinylation of proteins localized at the synaptic surface, as previously described^52^. After washing and scraping the neurons, 150 µg of lysate per condition were incubated overnight with 40µl of PierceTM Streptavidin Magnetic Beads slurry (Thermo Fisher Scientific 88817) at 4°C with gentle rotation. The next day, bead were washed as described before^52^ (2 × 1 mL RIPA lysis buffer, 1 × 1 mL of 1M KCl, 1x 1 mL of 0.1 M Na_2_CO_3_, 1 × 1 mL of 2 M urea in 10 mM Tris-HCl (pH 8.0), and again with 2 × 1 mL RIPA lysis buffer). Finally, to elute biotinylated proteins, beads were boiled for 10 min in 25 µl of 3x protein loading buffer supplemented with 20 mM DTT and 2 mM Biotin (Sigma, B4501). The streptavidin eluate was collected and run on an 10% SDS-PAGE gel and 0.22 µm nitrocellulose membranes were immunoblotted and developed as described below.

For the western blotting visualization of whole lysates, these were combined with 4x SDS protein loading buffer supplemented with 40 mM dithiothreitol DTT (Sigma, D9779), run on an 8% SDS-PAGE gel and transferred to a 0.22 µm nitrocellulose membrane (AmershamTM Protran, G10600080). For both western blots of whole lysates and biotinylated proteins, membranes were blocked with 10% Milk (Merck Milipore, 70166) in TBS-T (0.2% Tween-20 in Tris-buffered saline) at room temperature for 1 hour, then incubated with primary antibodies at 4°C overnight in gentle agitation. Anti-Lgi1/EPT (abcam 30868, 1:500 dilution), anti-GluA1 (UC Davis/NIH NeuroMab Facility, 75-327, 1:500 dilution) anti-V5 (Invitrogen R96025, 1:2000 dilution) and anti-Beta Actin (Thermo Fisher Scientific PA5-85271, 1:5000 dilution) antibodies were diluted in 10% milk. The following day, membranes were washed with 1x TBS-T four times for 10 min each time and probed with Goat Anti-Rabbit IgG (H + L)-HRP Conjugate or Goat Anti-Mouse IgG (H + L)-HRP secondary antibodies (BioRad 1706515 and 1706516, 1:5000 dilution in 10% milk), then washed with 1x TBS-T four times for 10 min each time and finally developed with Clarity or Clarity Max ECL Western Blotting Substrates (BioRad 1705060 and 1705062) using for imaging one ChemiDocTM Touch Imaging System (BioRad laboratories). For checking or to visualize the global biotinylation reaction, the membrane was blocked with 3% w/v BSA in TBS-T (0.2% Tween-20 in Tris-buffered saline) at 4°C overnight and incubated with Pierce™ High Sensitivity Streptavidin-HRP (Thermo Fisher Scientific 21130, 1:5000 dilution in 3% w/v BSA in TBS-T) at room temperature for 1 hr, then washed with TBS-T 3-4 times for 10 min each time and developed as described above. Uncropped blots corresponding to the blots shown in Figure 4 can be found in Supp. Figure S10.

#### Patient-derived polyclonal LGI1 auto-antibodies preparation and use

LGI1 IgG was purified from plasma exchange material of 3 patients suffering from LGI1 encephalitis, obtaining enriched samples with high titer (1:100) anti-LGI1 antibodies. These three samples were mixed as described previously^20^, and adjusted to a final concentration of 5 mg/ml in 0.9% NaCl for long term storage. Control IgGs, with a final concentration of 5 mg/ml, were obtained from a single donor who was clinically confirmed to not present any CNS disorder and who did not have detectable abnormal antibodies, as done previously^20,89^. All human subjects provided informed consent for use of plasma exchange material and use of human material was approved by the local ethics committee of Jena University Hospital (license # 2019-1415-Material).

Purified antibodies were added to primary rat hippocampal cultures at final concentrations of 100 µg/ml seven days before recordings. To avoid excessive usage of antibodies, a 10mm cloning cylinder was placed on top of the coverslip using vacuum grease, allowing to treat neurons contained inside the cylinder in a reduced volume of 120µl. One day before the experiment, half of the medium (60 µl) was removed from the media contained in the cylinder and purified polyclonal antibodies were added to the remaining medium (final concentration 100 µg/ml). Purified control antibodies from donors without LGI1-encephalitis were applied identically to the respective autoantibody groups. At the moment of the experiment, neurons were washed in Tyrode’s buffer and imaged as described above.

## Acknowledgments

We would like to thank all members of the de Juan-Sanz laboratory for insightful discussions and comments. This work was made possible by the Paris Brain Institute Diane Barriere Chair in Synaptic Bioenergetics awarded to Jaime de Juan-Sanz. Our funding sources are an ERC Starting Grant SynaptoEnergy (European Research Council, ERC-StG-852873), 2019 ATIP-Avenir Grant (CNRS, Inserm) and a Big Brain Theory Grant (ICM) awarded to J.d.J-S. C.P-C. is the recipient of a postdoctoral fellowship from the Fundacion Martin Escudero. J.d.J-S is a permanent CNRS researcher and a FENS-Kavli Scholar. This work was supported by an ERC Consolidator Grant (ERC-CoG-865634) to S.H and the German Research Foundation (FOR3004– SYNABS; [HA6386/10-2 to S.H.] and [GE2519/8-2 to C.G.]). MBH and ASA thank the generous support from NIH NINDS 5R01NS112365 (MBH) the NSF (CAREER 1750199) and the Neukom Institute (Dartmouth).

## Author contributions

U.C. and L.C-R. contributed equally to this work. Conceptualization, U.C., L.C-R. and J.d.J-S.; methodology, U.C., L.C-R., C.P-C. and J.d.J-S.; investigation, U.C., L.C-R., C.P-C. A.A.A, M.B.H. and J.d.J-S.; funding acquisition, project administration, and supervision, J.d.J-S.; providing essential materials; A.A., K.P., K.I., M.Y., C.G., S.H., A.R-J.; writing – original draft, U.C., L.C-R. and J.d.J-S.; writing – review & editing, U.C., L.C-R., C.P-C., A.A.A, M.B.H. and J.d.J-S

---

**Supplementary figure 1, related to main figure 1.**
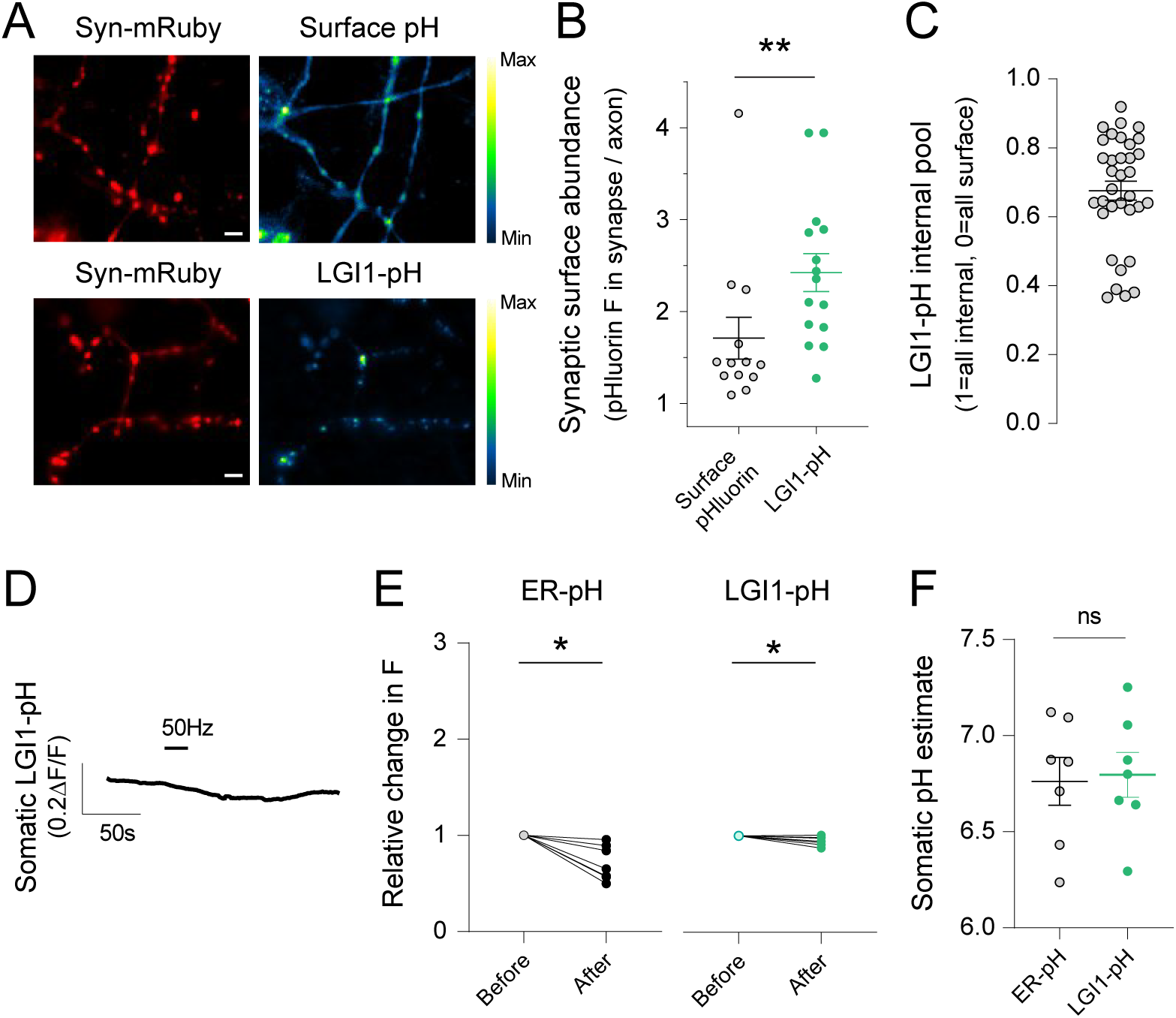
**(A)** Example images of surface pHluorin signals in axons (upper left) co-expressing synapsin-mRuby (upper left, colored in red) and LGI1 pHluorin signal in presynaptic boutons identified by synapsin-mRuby expression (lower left, colored in red). pHluorin signal shown as pseudocolor; scale shows low to high intensity. **(B)** Quantification of pHluorin abundance in synaptic surface of surface pHluorin versus LGI1-pHluorin expressing neurons. Data are mean ± SEM. Surface pH; n=13, 1.71 ± 0.23; LGI1-pH, n=15, 2.43 ± 0.21. Mann-Whitney test, **p<0.01. **(C)** Average internal pool of LGI1-pH, calculated using MES pH5.5 and NH_4_Cl pH 7.4 as shown in STAR methods. Bars are SEM. n=34, mean ± SEM, 0.68 ± 0.03. **(D)** Example trace of somatic LGI1-pH response to 1000AP 50Hz. **(E)** Relative fluorescence change in the soma before and after 1000AP 50Hz electrical stimulation in ER-pH (n=7) or LGI1-pH (n=7) expressing neurons. Wilcoxon matched-pairs signed rank test, *p<0.05. **(F)** Estimates of pH in somas of neurons expressing ER-pH or LGI1-pH. ER-pH, n=7, mean ± SEM, 6.8 ± 0.1 and LGI1-pH, n=7, mean ± SEM, 6.8 ± 0.1. Mann-Whitney test, ns.

**Supplementary figure 2, related to main figure 2.**
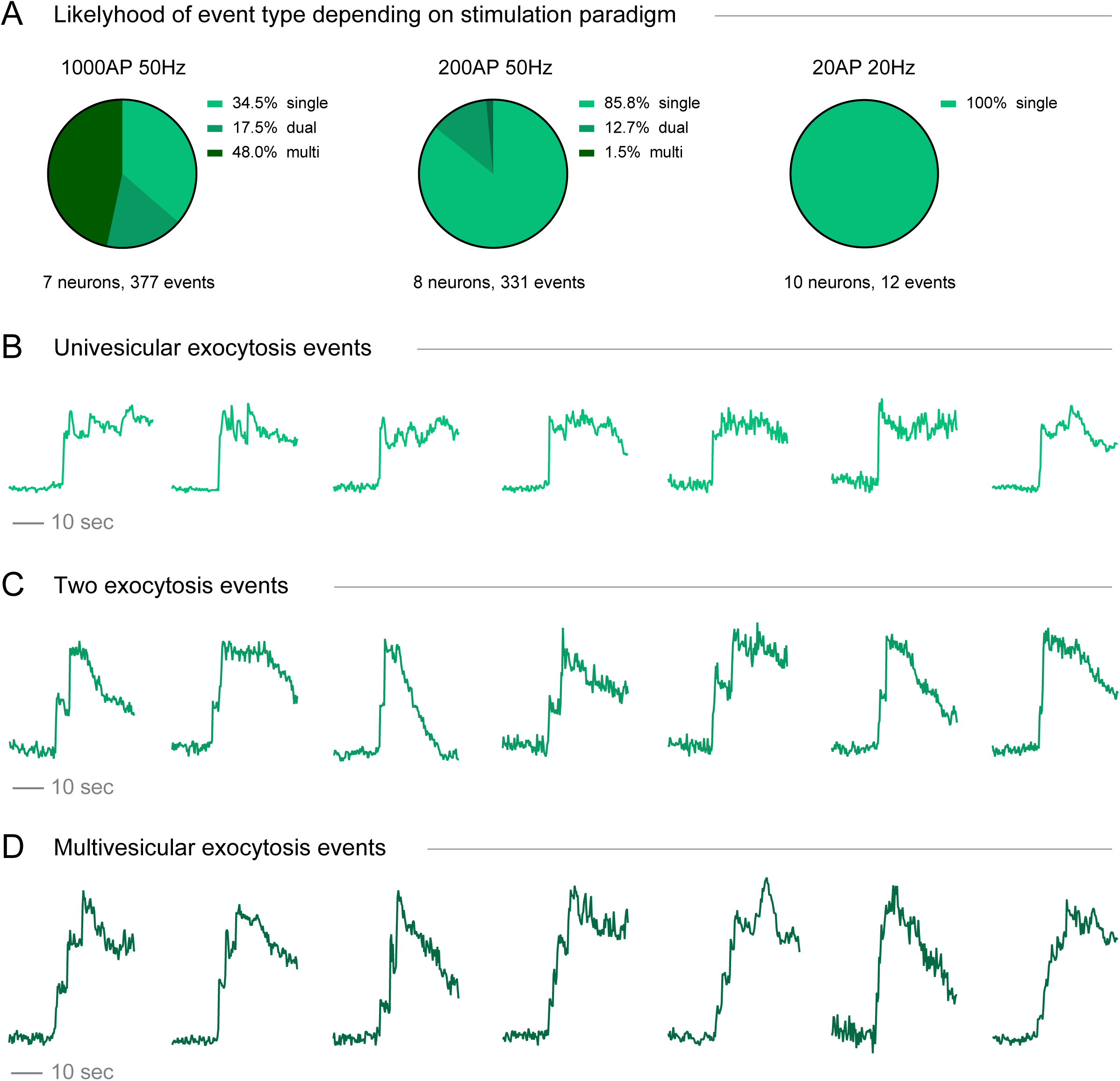
**(A)** Pie charts illustrating the distribution of LGI1-pH exocytosis event types based on the stimulation paradigm. For 1000AP 50Hz (n=377 events, 7 neurons) 34.5% events are univesicular, 17.5% present two events and 48.0% present multiple events. For 200AP 50Hz (n=331 events, 8 neurons), 85.8% exocytosis events are univesicular, 12.7% present two events and 1.5% present multiple events at. For 20AP 20Hz (n=12 events, 10 neurons), 100% events are univesicular. **(B)** Example traces for LGI1-pH univesicular exocytosis events during 1000AP 50Hz, normalized to the maximum peak response in each trace. **(C)** LGI1-pH example traces for exocytosis responses presenting two events during 1000AP 50Hz, normalized to the maximum peak response in each trace. **(D)** Example traces for LGI1-pH multivesicular exocytosis events at 1000AP 50Hz, normalized to the maximum peak response in each trace.

**Supplementary figure 3, related to main figure 3.**
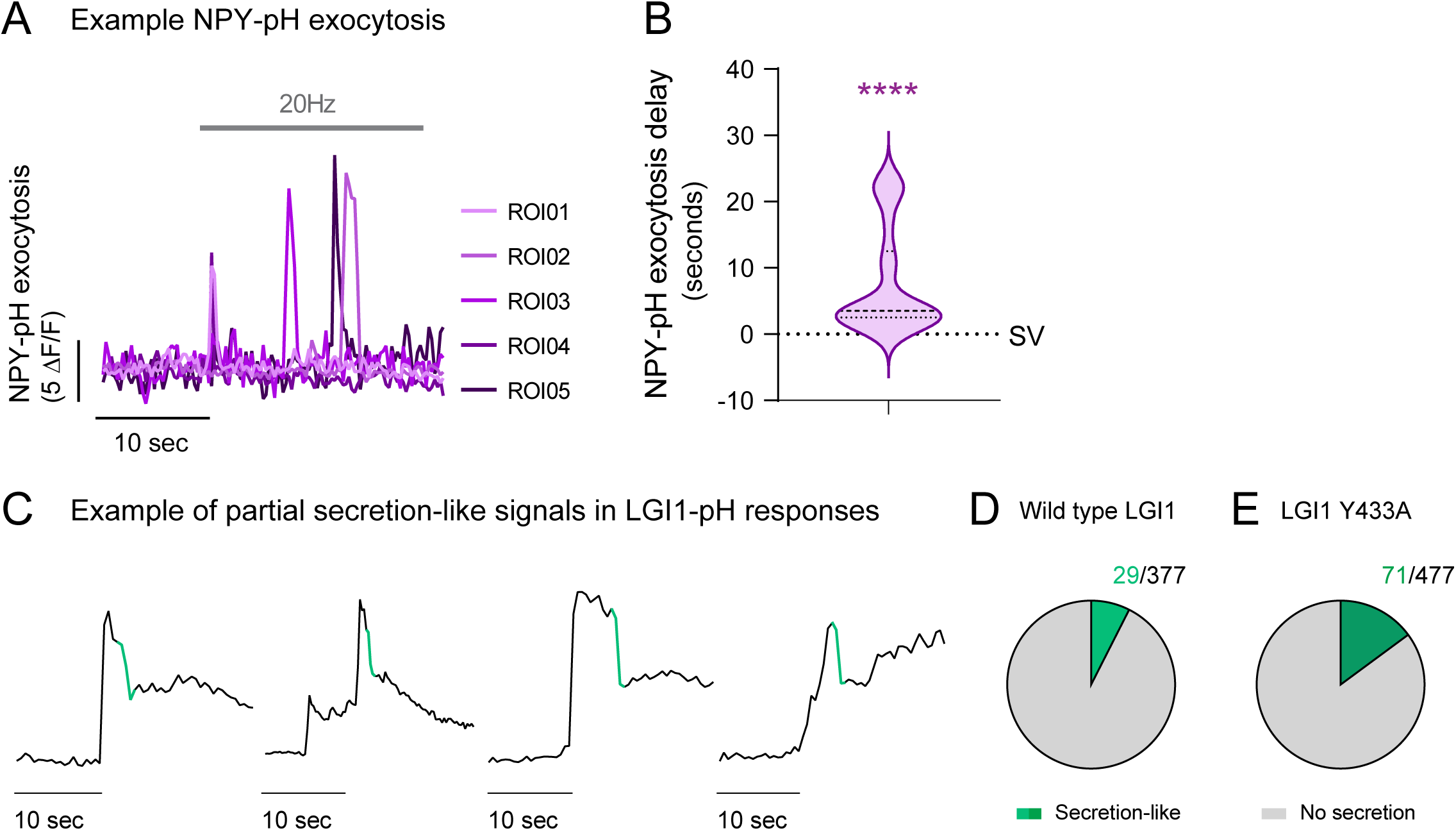
**(A)** NPY-pH example traces of secretion-like exocytosis obtained from a presynaptic arborization of a single neuron during a single 400AP, 20Hz stimulation. ROI means region of interest, which corresponds to a single bouton identified by synapsin-mRuby expression in the same neuron. **(B)** Quantification of delay in exocytosis of NPY-pH (n=39 boutons, 4 neurons) versus vGlut-pH (n=1214 boutons, 11 neurons; note that data is already shown in Figure 2). Mean ± SEM, 0.38 ± 0.01 seconds for vGlut-pH and 7.76 ± 1.28 seconds for NPY-pH. Mann-Whitney test; ****p<0.0001. **(C)** LGI1-pH example traces of sharp decreases in fluorescence compatible with partial secretion-like events. Each response was normalized to the maximum peak response in each trace shown. **(D)** Pie chart showing the identification of 29 secretion-like exocytosis events out of 377 events (7.7%) for wild type LGI1-pH (n=7 neurons). **(E)** Pie chart for LGI1^Y433A^-pH 71 secretion-like events out of 477 (14.9%; n=11 neurons).

**Supplementary figure 4, related to main figure 3.**
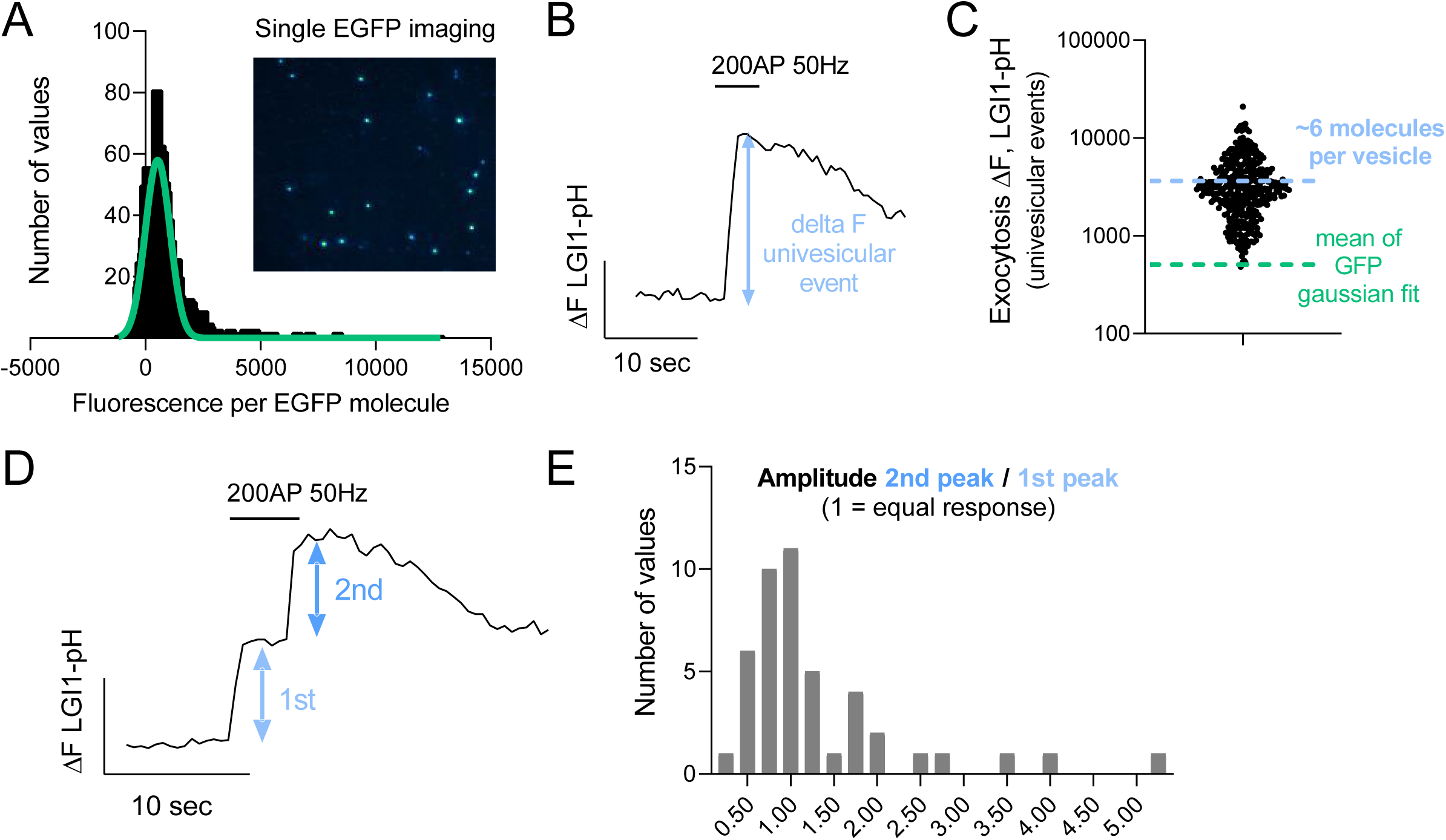
**(A)** Fluorescence distribution of 1727 single discrete EGFP bright spots. Bin size = 50. Data distribution was fitted to a single Gaussian distribution to obtain the equivalent fluorescence to a single EGFP molecule. Image shows an example of purified single EGFP molecules. **(B)** LGI1-pH example trace of a univesicular exocytosis event during 200AP 50Hz, imaged in the same conditions as the single EGFP imaging. Changes in F (ΔF) are quantified in each univesicular event as the one shown. **(C)** Individual ΔF univesicular responses, whose median shows an estimate of ∼6 LGI1-pH molecules exocytosis in these events (blue dashed line). Green dashed line shows the median fluorescence of a single EGFP molecule calculated as in A. **(D)** Example LGI1-pH trace of two exocytosis events during 200AP 50Hz. **(E)** Histogram showing the distribution of the relative amplitudes of the first and second event in those boutons that presented two exocytosis events. Values per bouton are calculated as ΔF(1)/ΔF(2), thus 1 indicates an equal change in ΔF in first and second events. Bin size = 0.5.

**Supplementary figure 5, related to main figure 4.**
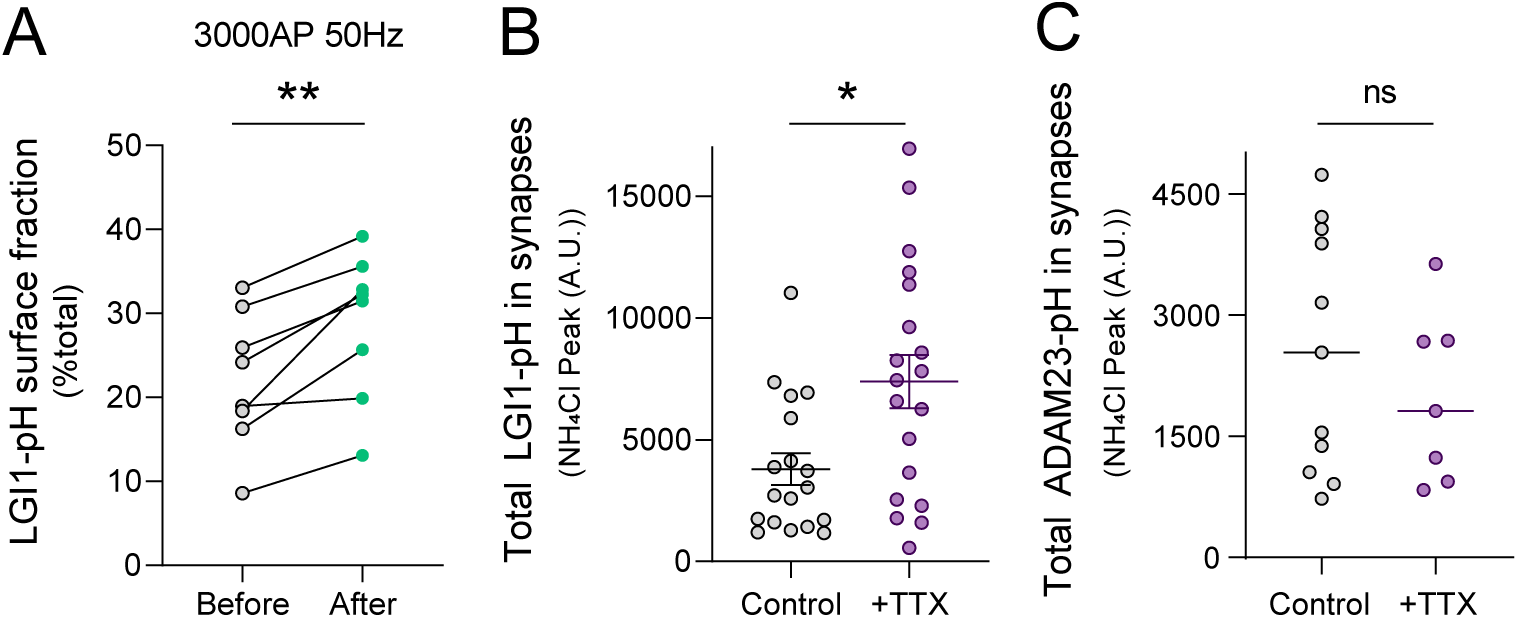
**(A)** LGI1-pH surface fraction before and after electrical stimulation of 3000AP 50Hz in individual neurons (n=8 neurons). Wilcoxon matched-pairs signed rank test, **p<0.01. **(B)** Total LGI1-pH abundance in synapses in control versus TTX treated neurons, measured as peak response in NH_4_Cl pH 7.4. Control, n=18, Mean ± SEM 3789 ± 653 and TTX, n=19, Mean ± SEM 7385 ± 1094. Mann-Whitney test, *p<0.05. **(C)** Total ADAM23-pH abundance in synapses in control versus TTX treated neurons, measured as peak response in NH_4_Cl pH 7.4. Control, n=11, Mean ± SEM 2566 ± 454 and TTX, n=7, Mean ± SEM 1974 ± 399.5. Mann-Whitney test, n.s.

**Supplementary figure 6, related to main figure 5.**
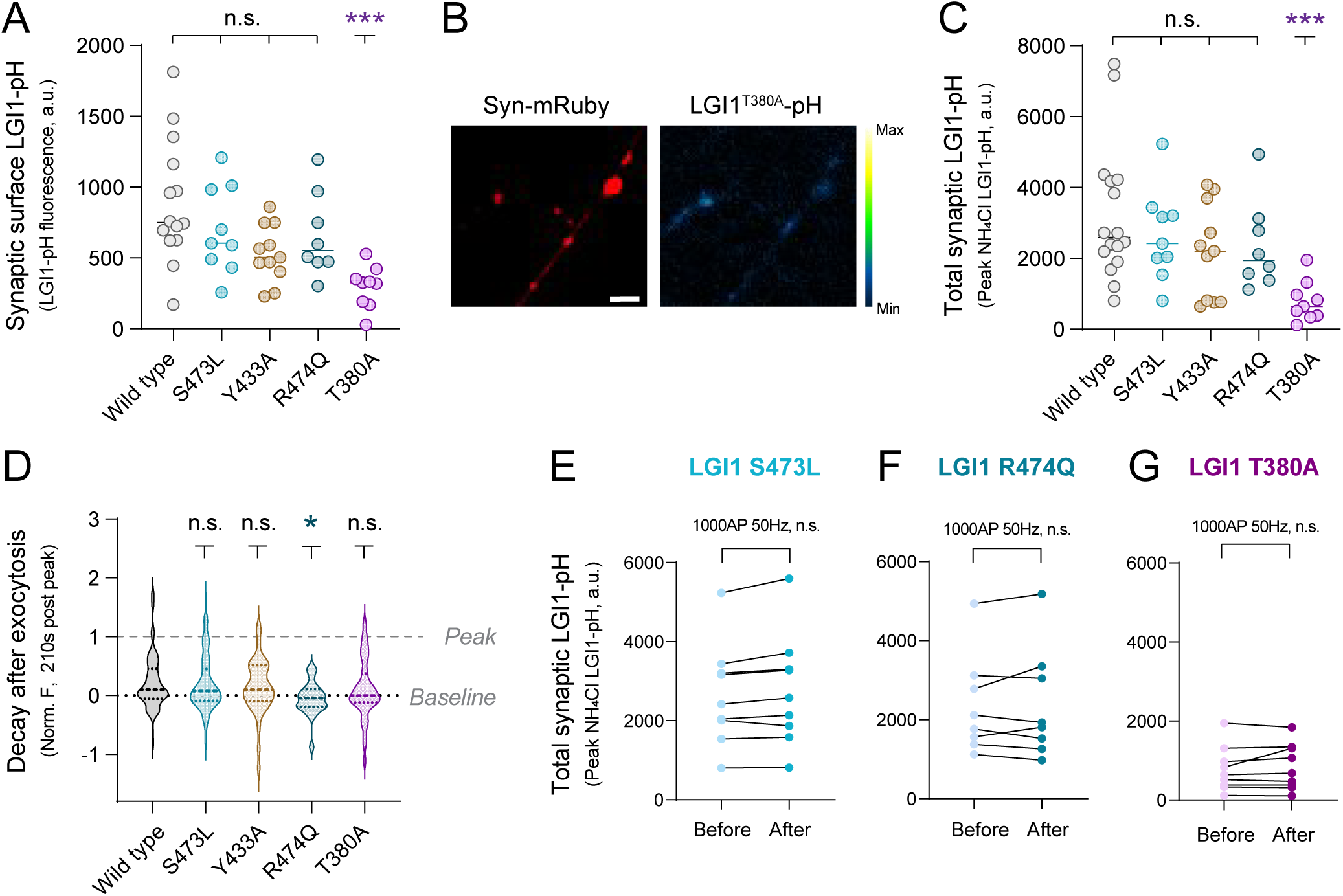
**(A)** Synaptic surface LGI1 at rest in pHluorin-tagged LGI1 constructs: WT, S473L, Y433A, R474Q and T380A. Data are mean ± SEM in a.u., WT, n=14, 895.3 ± 116.4; S473L, n=9, 697.5 ± 103.2; Y433A, n=11, 530.7 ± 60.6; R474Q, n=8, 658.3 ± 104.7; T380A, n=9, 296.9 ± 49.1. Dunn’s multiple comparisons test of WT peak response against S473L, Y433A or R474Q showed no statistical difference (n.s.). WT vs. T380A, ***p<0.001. **(B)** LGI1^T380A^-pH signal in presynaptic boutons (right, pseudocolor) identified by synapsin-mRuby expression (left, colored in red). Scale bar = 4.8 μm. **(C)** Total abundance in synapses of LGI1-pH constructs: WT, S473L, Y433A, R474Q or T380A, measured as peak response in NH_4_Cl pH 7.4. Data are mean ± SEM in a.u., WT, n=16, 3230 ± 479; S473L, n=9, 2649 ± 430; Y433A, n=11, 2188 ± 400; R474Q, n=8, 2350 ± 442; T380A, n=9, 784 ± 189. Dunn’s multiple comparisons test of WT abundance against S473L, Y433A or R474Q showed no statistical difference (n.s.). WT vs. T380A ***p<0.001. **(D)** Quantification of fluorescence decay in individual boutons 210 seconds after exocytosis peak for LGI1-pH constructs: WT, S473L, Y433A, R474Q or T380A. Data are normalized per bouton to the maximum fluorescence of the peak response in that bouton. Data are mean ± SEM, WT, n= 130, 0.22 ± 0.04; S473L, n= 91, 0.20 ± 0.05; Y433A, n=42, 0.14 ± 0.07; R474Q, n= 24, -0.05 ± 0.06; T380A, n= 48, 0.10 ± 0.07. Dunn’s multiple comparisons test of WT decay against S473L, Y433A or R474Q showed no statistical difference (n.s.). *p<0.05; WT vs. T380A n.s. **(E-G)** Quantification of total levels of pHluorin-tagged LGI1 constructs: S473L (E; n=9), R474Q (F; n=8) and T380A (G; n=9), before and after 1000AP 50Hz stimulation. Wilcoxon matched-pairs signed rank tests, n.s.

**Supplementary figure 7, related to main figure 5.**
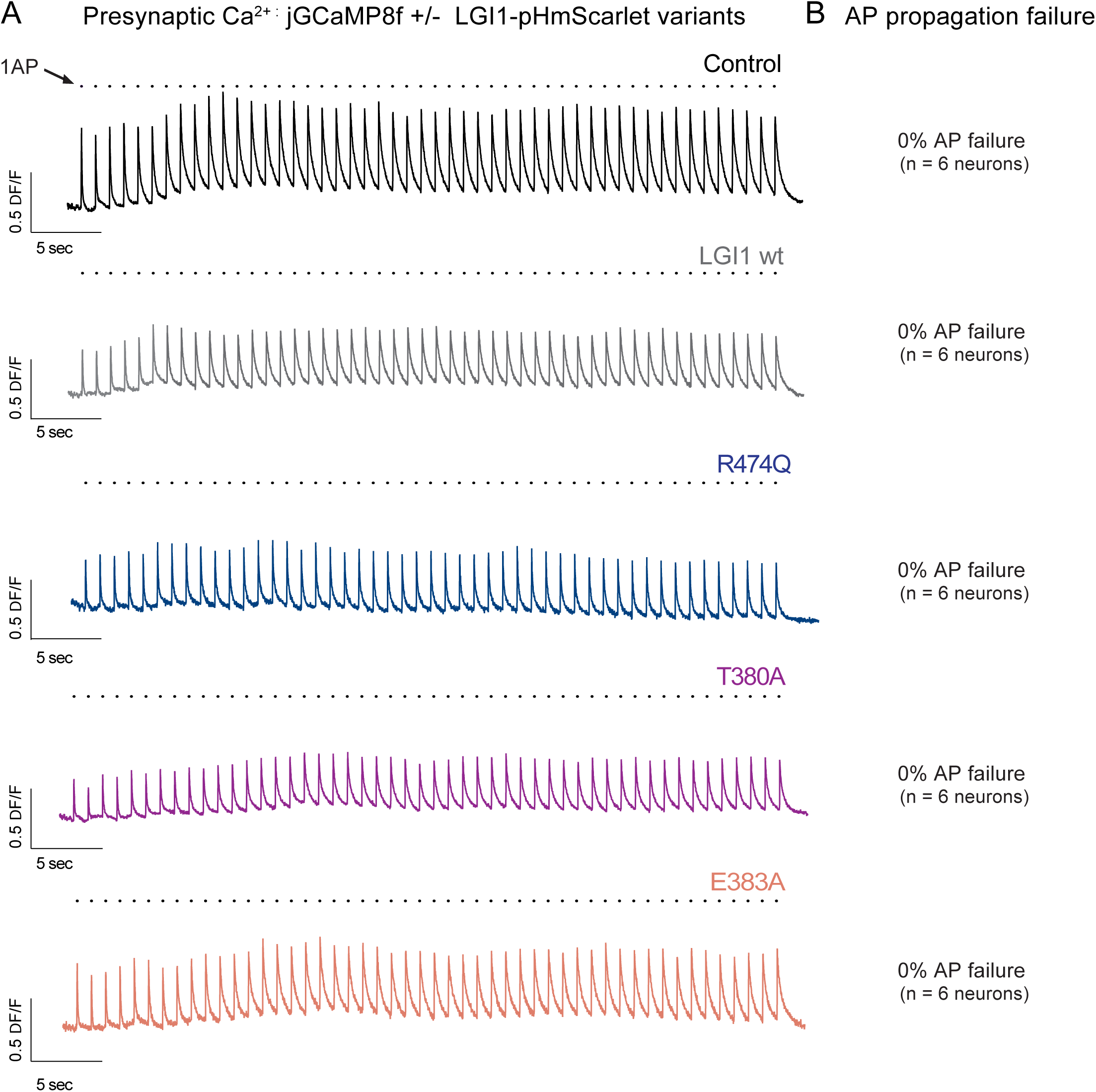
**(A)** Example presynaptic calcium transients from individual neurons stimulated to fire one action potential every second during 50 seconds. Responses are obtained from a single round of stimulation, as there is sufficient signal to noise to estimate possible failures. Neurons expressed only jGCaMP8f (top, black traces), or jGCaMP8f with pHmScarlet-tagged LGI1 construct WT, R474Q, T380A and E383A, shown as gray, blue, purple and orange traces, respectively. Note every time a single stimulation is induced, a robust response is found at the presynapse of single neurons. **(B)** AP propagation failure quantified as percentage: jGCaMP8f 0% (300 AP, 6 neurons); jGCaMP8f + LGI1-pHmScarlet 0% (300 AP, 6 neurons); jGCaMP8f + LGI1^R474Q^-pHmScarlet 0% (294 AP, 6 neurons); jGCaMP8f + LGI1^T380A^-pHmScarlet 0% (300 AP, 6 neurons); jGCaMP8f + LGI1^E383A^-pHmScarlet 0% (300 AP, 6 neurons).

**Supplementary figure 8.**
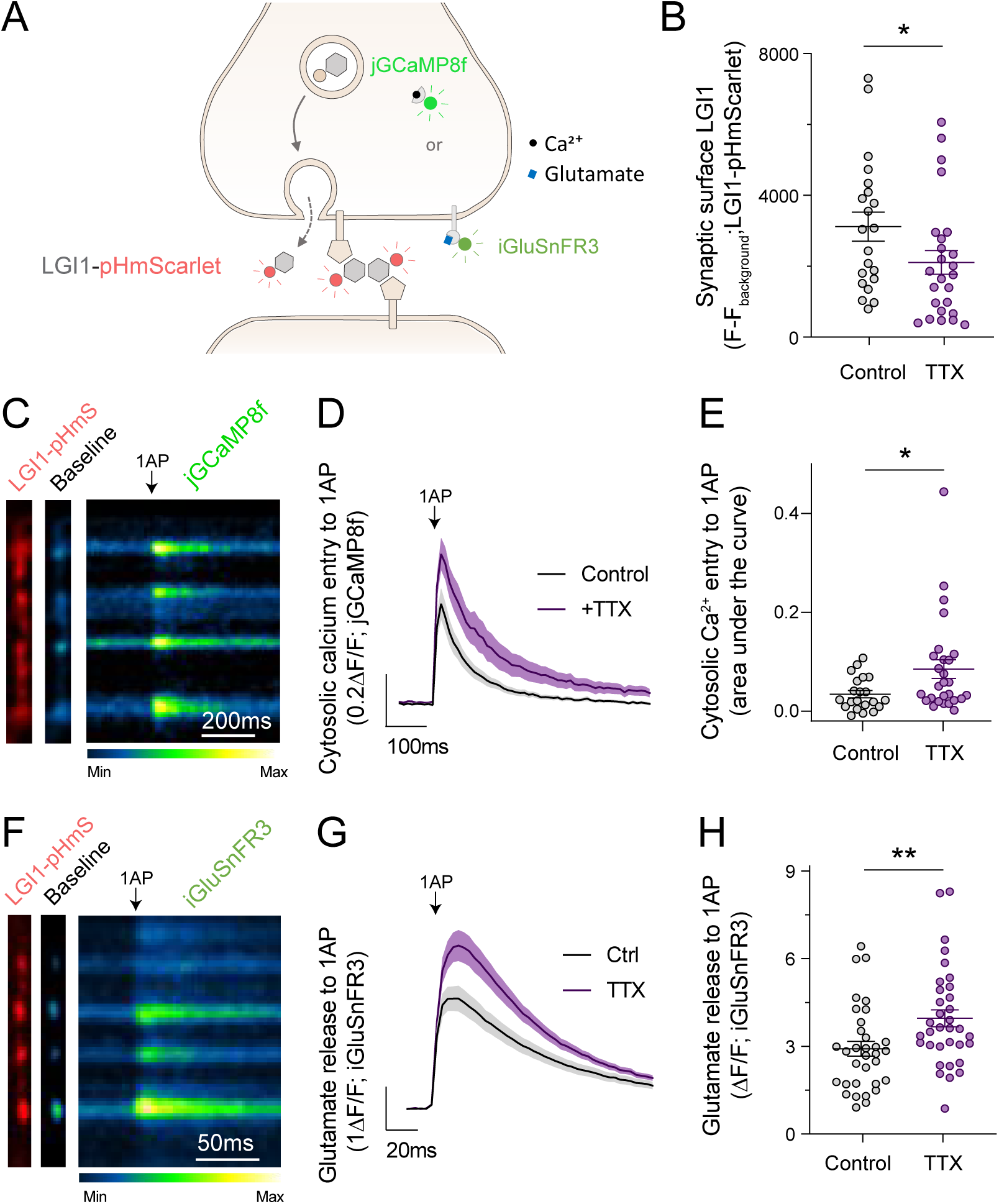
**(A)** Diagram showing LGI1-pHmScarlet (LGI1-pHmS), which is only bright when exposed to the extracellular media. **(B)** Effect of TTX on the LGI1-pHmScarlet abundance at the synaptic surface. Data are Mean ± SEM. Control, n=21, 3116 ± 408; TTX, n=25, 2108 ± 334. Mann-Whitney, *p<0.05. **(C)** LGI1-pHmScarlet signals in boutons (left, colored in red) co-expressing jGCaMP8f at rest (middle image) and during a 1AP stimulus (displayed as a kymograph, right; pseudocolor scale shows low to high intensity). Scale bar 200ms. **(D)** Average traces showing cytosolic calcium entry to 1AP stimulus in control versus TTX treated neurons. n=21 and n=27 cells, respectively. Error bars are SEM. **(E)** Quantification of peak cytosolic calcium entry responses to 1AP stimulus in control versus TTX treated neurons. Data are mean ± SEM. Control n=21, 0.03 ± 0.01 ΔF/F; TTX n=27, 0.09 ± 0.02 ΔF/F. Mann-Whitney test, *p<0.05. **(F)** LGI1-pHmScarlet signals in boutons (left, colored in red) co-expressing iGluSnFR3 at rest (middle image) and during a 1AP stimulus (displayed as a kymograph, right; pseudocolor scale shows low to high intensity). Scale bar, 50ms. **(G)** Average traces showing glutamate release to 1AP stimulus in control versus TTX treated neurons; n=33 and n=34 neurons, respectively. Bars are SEM. **(H)** Quantification of peak glutamate release to 1AP stimulus in control versus TTX treated neurons. Data are mean ± SEM. Control n=33, 2.94 ± 0.25; TTX n=34, 3.96 ± 0.29. Mann-Whitney test, **p<0.01.

**Supplementary figure 9, related to main figures 3, 6 and 7.**
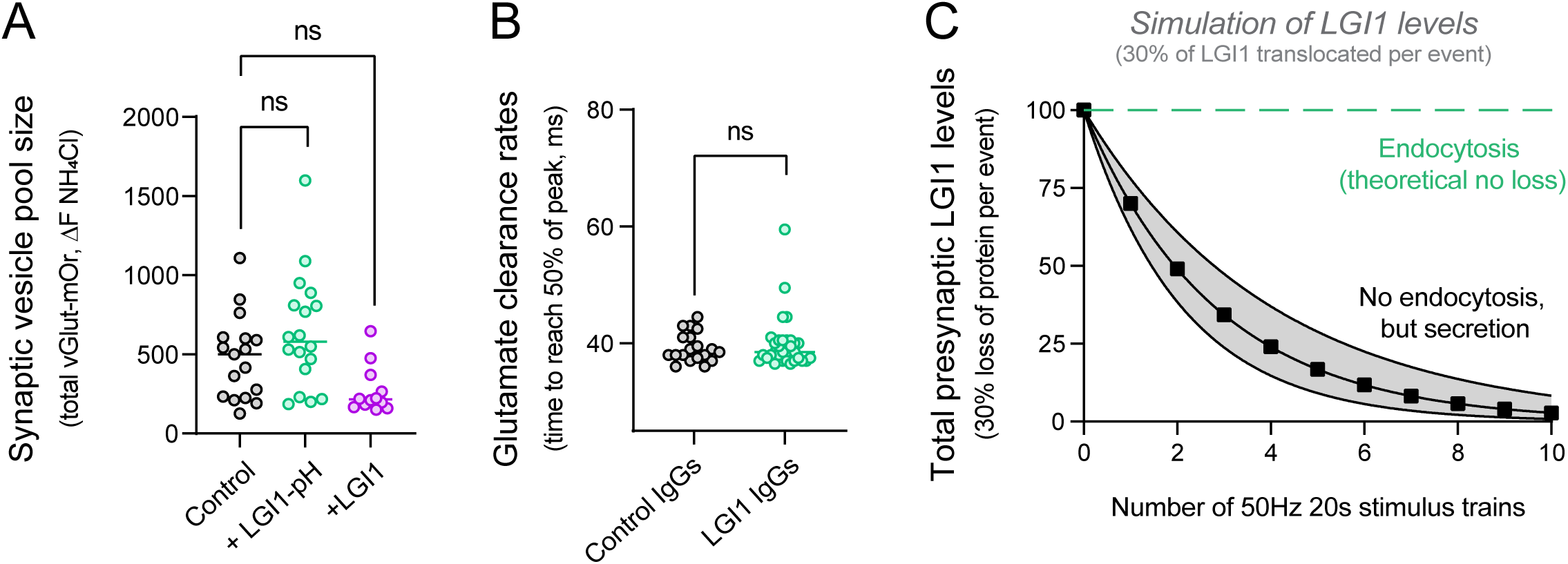
**(A)** Total pool of synaptic vesicles in control versus LGI1-pH or untagged LGI1 overexpression, measured as vGlut-mOrange-2 peak response in NH_4_Cl pH 7.4. Data are mean ± SEM, Control, n=17, 477.8 ± 64.3; LGI1-pH, n=18, 635.7 ± 85.2 for; LGI1, n=12, 272.0 ± 43.8. Dunn’s multiple comparisons test of the abundance of control vs. LGI1-pH or untagged LGI1 showed no statistical difference (n.s.) **(B)** Presynaptic glutamate clearance rates were measured as the time to reach half of the peak response per neuron. Clearance rates were measured in neurons expressing LGI1-pH and treated with either IgGs from control or anti-LGI1 IgGs from limbic encephalitis patients. Data are mean ± SEM. Control IgGs n=19, 39.3 ± 0.6; LGI1 IgGs n=32, 39.9 ± 0.8 for LGI1 IgGs. t-test, n.s. **(C)** Simulation of total LGI1 levels in the presynapse during 50Hz 20s stimulus rounds, assuming either no loss of protein through endocytosis or a 30% loss of protein through secretion per stimulation round.

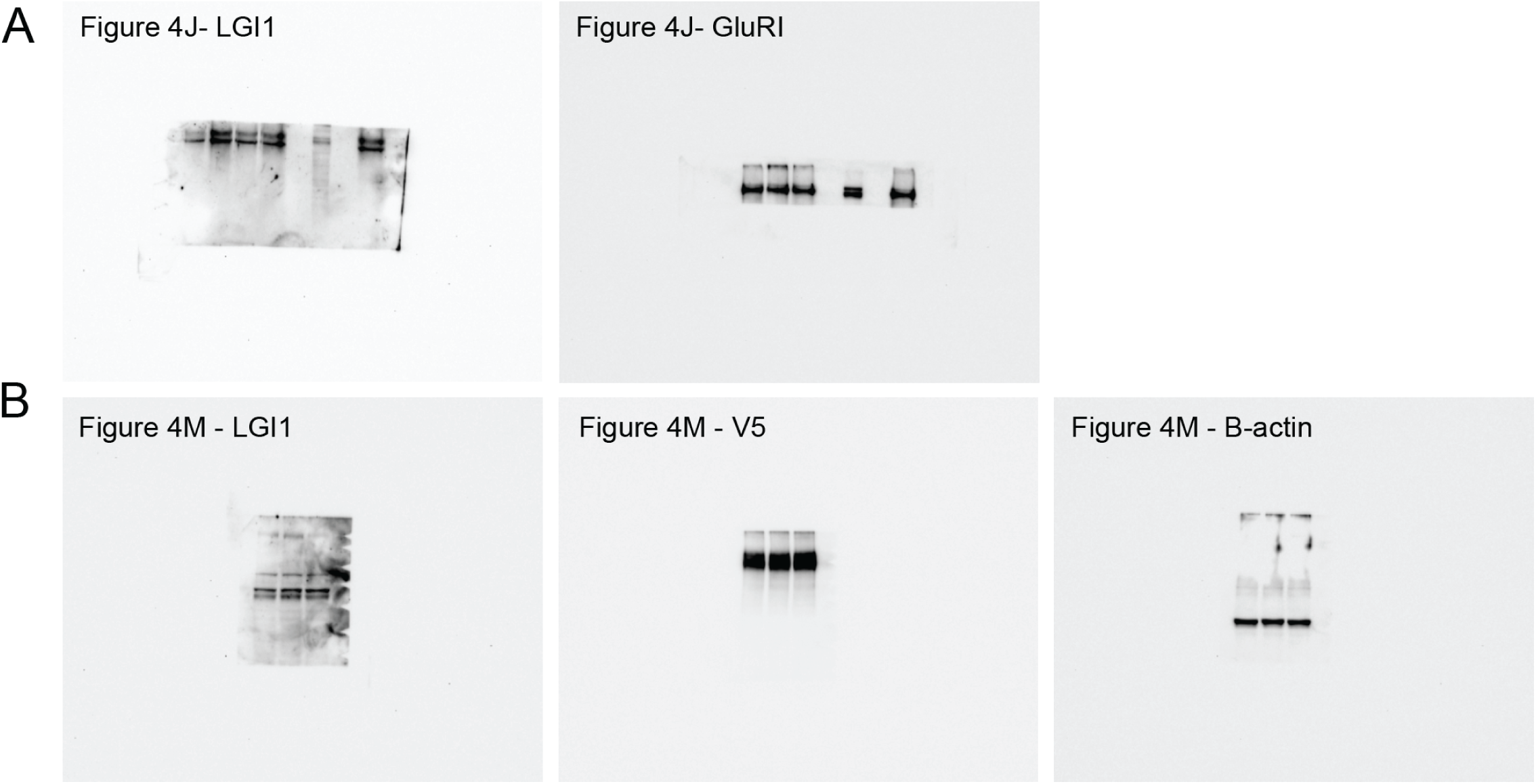

